# VIBRANT: Automated recovery, annotation and curation of microbial viruses, and evaluation of virome function from genomic sequences

**DOI:** 10.1101/855387

**Authors:** Kristopher Kieft, Zhichao Zhou, Karthik Anantharaman

## Abstract

**Background:** Viruses are central to microbial community structure in all environments. The ability to generate large metagenomic assemblies of mixed microbial and viral sequences provides the opportunity to tease apart complex microbiome dynamics, but these analyses are currently limited by the tools available for analyses of viral genomes and assessing their metabolic impacts on microbiomes.

**Design:** Here we present VIBRANT, the first method to utilize a hybrid machine learning and protein similarity approach that is not reliant on sequence features for automated recovery and annotation of viruses, determination of genome quality and completeness, and characterization of virome function from metagenomic assemblies. VIBRANT uses neural networks of protein signatures and a novel v-score metric that circumvents traditional boundaries to maximize identification of lytic viral genomes and integrated proviruses, including highly diverse viruses. VIBRANT highlights viral auxiliary metabolic genes and metabolic pathways, thereby serving as a user-friendly platform for evaluating virome function. VIBRANT was trained and validated on reference virus datasets as well as microbiome and virome data.

**Results:** VIBRANT showed superior performance in recovering higher quality viruses and concurrently reduced the false identification of non-viral genome fragments in comparison to other virus identification programs, specifically VirSorter and VirFinder. When applied to 120,834 metagenomically derived viral sequences representing several human and natural environments, VIBRANT recovered an average of 94.5% of the viruses, whereas VirFinder and VirSorter achieved less powerful performance, averaging 48.1% and 56.0%, respectively. Similarly, VIBRANT identified more total viral sequence and proteins when applied to real metagenomes. When compared to PHASTER and Prophage Hunter for the ability to extract integrated provirus regions from host scaffolds, VIBRANT performed comparably and even identified proviruses that the other programs did not. To demonstrate applications of VIBRANT, we studied viromes associated with Crohn’s Disease to show that specific viral groups, namely Enterobacteriales-like viruses, as well as putative dysbiosis associated viral proteins are more abundant compared to healthy individuals, providing a possible viral link to maintenance of diseased states.

**Conclusions:** The ability to accurately recover viruses and explore viral impacts on microbial community metabolism will greatly advance our understanding of microbiomes, host-microbe interactions and ecosystem dynamics.

## Background

Viruses that infect bacteria and archaea are incredibly abundant, and outnumber their hosts in most environments (*1–3*). Viruses are commonly considered non-living entities and are obligate intracellular pathogenic genetic elements capable of reprogramming host cellular metabolic states during infection. They are also highly active and cause the lysis of 20-40% of microorganisms in diverse environments every day (*4, 5*). Due to their abundance and widespread activity, viruses are vital to microbial communities as they drive cycling of essential nutrients such as carbon, nitrogen, phosphorus and sulfur (*6–10*). In human systems, viruses have been implicated in contributing to dysbiosis that can lead to various diseases, such as inflammatory bowel diseases, or even have a symbiotic role with the immune system (*11–13*).

It is estimated that viral diversity exceeds that of living organisms, and therefore harbors enormous potential for diverse genomic content, arrangement and encoded functions (*14*). Accordingly, there is substantial interest in “mining” viral sequences for novel anti-microbial drug candidates, enzymes for biotechnological applications, and for bioremediation efforts (*15–19*). Moreover, viruses have a unique capability to rapidly evolve genes via high mutation rates and act as intermediate carriers to transfer these genes to their hosts and subsequently to the surrounding communities (*20–22*).

Our understanding of the diversity of viruses continues to expand with the discovery of novel viral lineages within the last decade. One of the most striking examples is the characterization of crAssphage, an extraordinarily abundant virus infecting *Bacteroides* species within the human gut that went unnoticed for years due to its lack of homology with known viral sequences (*23*). Moreover, the discovery of megaphages, the largest known bacterial viruses infecting the human gut bacteria *Prevotella*, has pushed the boundaries on the coding capacity of viruses (*24, 25*). In the oceans, a newly discovered lineage of *Vibrio*-infecting non-tailed viruses, generally considered unconventional since most known bacterial viruses are tailed, fueled the notion that current viral recovery methods are skewing our understanding of viruses in the environment (*26*). Taken together, this highlights that estimates of viral diversity are biased towards tailed dsDNA viruses and are likely underrepresenting other families of viruses including those with ssDNA and RNA genomes (*27, 28*).

Recently it has been appreciated that viruses may directly link biogeochemical cycling of nutrients by specifically driving metabolic processes. For example, during infection viruses can acquire 40-90% of their required nutrients from the surrounding environment by taking over and subsequently directing host metabolism (*29–31*). To manipulate host metabolic frameworks some viruses have selectively “stolen” metabolic genes from their host. These host derived genes, collectively termed auxiliary metabolic genes (AMGs), are actively expressed during infection to provide viruses with fitness advantages (*32–35*). Viruses encoding AMGs have been found to be widespread in human and natural environments and implicated in manipulating several important nutrient cycles including carbon, nitrogen, phosphorus and sulfur (*36–40*). Identifying these genes and understanding the processes underpinning their function is pivotal for developing comprehensive models of the impacts of microbiomes and nutrient cycling.

Due to the difficulty of collecting virus-only samples as well as the need to integrate viruses into models of ecosystem function, it has become of great interest to determine which sequences within microbial communities are derived from viruses. Within the cellular fraction of a sample there can remain a large number of viruses for a variety of reasons. First, these viruses can exist as active intracellular infections, which may be the case for as many as 30% of all bacteria at any given time (*41*). Second, there may be particle-attached viruses resulting from viruses’ inherently “sticky” nature (*42*). Lastly, many viruses exist as “proviruses”, or viral genomes either integrated into that of their host or existing within the host as an episomal sequence. As such, it is crucial for the accurate evaluation of microbial community characteristics, structure and functions to be able to separate these viral sequences.

Multiple tools exist for the identification of viral sequences from mixed metagenomic assemblies. For several years VirSorter (*43*), which succeeded tools such as VIROME (*44*) and Metavir (*45*), has been the most widely used for its ability to accurately identify viral metagenomic fragments (scaffolds) from large metagenomic assemblies. VirSorter predominantly relies on database searches of predicted proteins, using both reference homology as well as probabilistic similarity, to compile metrics of enrichment of virus-like proteins and simultaneous depletion of other proteins. To do this it uses a virus-specific curated database as well as Pfam (*46*) for non-virus annotations, though it does not fully differentiate viral from non-viral Pfam annotations. It also incorporates signatures of viral genomes, such as encoding short genes or having low levels of strand switching between genes. VirSorter is also unique in its ability to use these annotation and sequence metrics to identify and extract integrated provirus regions from host scaffolds. After prediction of viral sequences, VirSorter labels viral scaffolds with one of three confidence levels: *categories* 1, 2 or 3. Categories 1 and 2 are generally considered accurate, but category 3 predictions are more likely to contain false identifications. While VirSorter is quite accurate, it likely underrepresents the diversity and abundance of viruses within metagenomic assemblies.

More recent tools have been developed to compete with the performance of VirSorter in order to expand our appreciation and understanding of viruses. VirFinder (*47*) was the first tool to implement machine learning and be completely independent of reference databases for predicting viral sequences which was a platform later implemented in PPR-Meta (*48*). VirFinder was built with the consideration that viruses tend to display distinctive patterns of 8-nucleotide frequencies (otherwise known as 8-mers), which was proposed despite the knowledge that viruses can share remarkably similar nucleotide patterns with their host (*49*). These 8-mer patterns were used to build a random forest machine learning model to quickly classify sequences as short as 500 bp without the need for gene prediction. VirFinder generates model-derived scores as well as probabilities of prediction accuracy, though it is up to the user to define the cutoffs which can ultimately lead to uncertainties in rates of false identification of viral sequences. VirFinder was shown to greatly improve the ability to recover viral sequences compared to VirSorter, but it also demonstrates substantial host and source environment biases in predicting diverse viruses. For example, VirFinder was able to recover viruses infecting Proteobacteria more readily than those infecting Firmicutes due to reference database-associated biases while training the machine learning model. Additional biases were also identified between different source environments, seen through the under-recovery of viruses from certain environments compared to others (*50*).

Additional recent tools have been developed that utilize slightly different methods for identifying viral scaffolds. MARVEL (*51*), for example, leverages annotation, sequence signatures (e.g., strand switching and gene density) and machine learning to identify viruses from metagenomic bins. MARVEL differs from VirSorter in that it only utilizes a single virus-specific database for annotation and also differs from VirFinder in that it does not use global nucleotide frequency patterns. However, MARVEL provides no consideration for integrated proviruses and is only suitable for identifying bacterial viruses from the order *Caudovirales* which substantially limits its ability to discover novel viruses. Another recently developed tool, VirMiner (*52*), is unique in that it functions to use metagenomic reads and associated assembly data to identify viruses and performs best for high abundance (i.e., high coverage when assembled) viruses. VirMiner is a web-based server that utilizes a hybrid approach of employing both homology-based searches to a virus-specific database as well as machine learning. VirMiner was found to have improved ability to recover viral scaffolds compared to both VirSorter and VirFinder but was concurrently much less accurate. Poor accuracy would lead to a skewed interpretation of virome function if the identified virome consisted of many non-viral sequences. This distinction is important because VirMiner employs functional characterization as well as determination of virus-host relationships.

Thus far, VirSorter remains the most efficient tool for identifying integrated proviruses within metagenomic assemblies. Other tools, predominantly PHASTER (*53*) and Prophage Hunter (*54*), are specialized in identifying integrated proviruses from whole genomes rather than scaffolds generated by metagenomic assemblies. Similar to VirSorter, these two provirus predictors rely on reference homology and viral sequence signatures with sliding windows to identify regions of a host genome that belong to a virus. Although they are useful for whole genomes, they lack the capability of identifying scaffolds belonging to lytic (i.e., non-integrated) viruses and perform slower for large datasets. In addition, both PHASTER and Prophage Hunter are exclusively available as web-based servers and offer no stand-alone command line tools.

Here we developed VIBRANT (Virus Identification By iteRative ANnoTation), a tool for automated recovery, annotation, and curation of both free and integrated viruses from metagenomic assemblies and genome sequences. VIBRANT is capable of identifying diverse dsDNA, ssDNA and RNA viruses infecting both bacteria and archaea, and to our knowledge has no evident environmental biases. VIBRANT uses neural networks of protein annotation signatures from non-reference-based similarity searches with Hidden Markov Models (HMMs) as well as a unique ‘v-score’ metric to maximize identification of diverse and novel viruses. After identifying viral scaffolds VIBRANT implements curation steps to validate predictions. VIBRANT additionally characterizes virome function by highlighting AMGs and assesses the metabolic pathways present in viral communities. All viral genomes, proteins, annotations and metabolic profiles are compiled into formats for user-friendly downstream analyses and visualization. When applied to reference viruses, non-reference virus datasets and various assembled metagenomes, VIBRANT outperformed both VirFinder and VirSorter in the ability to maximize virus recovery and minimize false discovery. When compared to PHASTER and Prophage Hunter for the ability to extract integrated provirus regions from host scaffolds, VIBRANT performed comparably and even identified proviruses that the other programs did not. VIBRANT was also used to identify differences in metabolic capabilities between viruses originating from various environments. When applied to three separate cohorts of individuals with Crohn’s Disease, VIBRANT was able to identify both differentially abundant viral groups compared to healthy controls as well as virally encoded genes putatively influencing a diseased state. VIBRANT is freely available for download at https://github.com/AnantharamanLab/VIBRANT. VIBRANT is also available as a user-friendly, web-based application through the CyVerse Discovery Environment at https://de.cyverse.org/de/?type=apps&app-id=c2864d3c-fd03-11e9-9cf4-008cfa5ae621&system-id=de (*55*).

## Results

VIBRANT was built to extract and analyze bacterial and archaeal viruses from assembled metagenomic and genome sequences, as well as provide a platform for characterizing metabolic proteins and functions in a comprehensive manner. The concept behind VIBRANT’s mechanism of virus identification stems from the understanding that arduous manual inspection of annotated genomic sequences produces the most dependable results. As such, the primary metrics used to inform validated curation standards and to train VIBRANT’s machine learning based neural network to identify viral sequences reflects human-guided intuition, though in a high-throughput automated fashion.

### Determination of v-score

We developed a unique ‘v-score’ metric as an approach for providing quantitative information to VIBRANT’s algorithm in order to assess the qualitative nature of annotation information. A v-score is a value assigned to each possible protein annotation that scores its association to viral genomes. V-score differs from the previously used “virus quotient” metric (*56, 57*) in that it does not take into account the annotation’s relatedness to bacteria or archaea. Not including significant similarity to non-viral genomes in the calculation of v-scores has important implications for this metric’s utility. Foremost is that annotations shared between viruses and their hosts, such as ribonucleotide reductases, will be assigned a v-score reflecting its association to viruses, not necessarily virus-specificity. Many genes are commonly associated with viruses and host organisms, but when encoded on viral genomes can be central to virus replication efficiency (e.g., ribonucleotide reductases (*58*)). Therefore, a metric representing virus-association rather than virus-specificity would be more appropriate in identifying if an unknown scaffold is viral or not. Secondly, this approach takes into account widespread horizontal gene transfer of host genes by viruses as well as the presence of AMGs.

### VIBRANT workflow

VIBRANT utilizes several annotation metrics in order to guide removal of non-viral sequences before curation of reliable viral scaffolds. The annotation metrics used are derived from HMM-based probabilistic searches of protein families from the Kyoto Encyclopedia of Genes and Genomes (KEGG) KoFam (*59, 60*), Pfam (*46*) and Virus Orthologous Group (VOG) (vogdb.org) databases. VIBRANT is not reliant on reference-based similarity and therefore accounts for the large diversity of viruses on Earth and their respective proteins. Consequently, widespread horizontal gene transfer, rapid mutation and the vast amount of novel sequences do not hinder VIBRANT’s ability to identify known and novel viruses. VIBRANT relies minimally on non-annotation features, such as rates of open reading frame strand switching, because these features were not as well conserved in genomic scaffolds in contrast to whole genomes.

VIBRANT’s workflow consists of four main steps (Figure 1A). Briefly, proteins (predicted or user input) are used by VIBRANT to first eliminate non-viral sequences by assessing non-viral annotation signatures derived from KEGG and Pfam HMM annotations. At this step potential host scaffolds are fragmented using sliding windows of KEGG v-scores in order to extract integrated provirus sequences. Following the elimination of most non-viral scaffolds and rough excision of provirus regions, proteins are annotated by VOG HMMs. Before analysis by the neural network machine learning model, any extracted putative provirus is trimmed to exclude any remaining non-viral sequences. Annotations from KEGG, Pfam and VOG are used to compile 27 metrics that are utilized by the neural network to predict viral sequences. These 27 metrics were found to be adequate for the separation of viral and non-viral scaffolds (Figure 1B). After prediction by the neural network a set of curation steps are implemented to filter the results in order to improve accuracy as well as recovery of viruses. Once viruses are identified VIBRANT automates the analysis of virome function by highlighting AMGs and assigning them to KEGG metabolic pathways. The genome quality (i.e., proxy of completeness) of identified viral scaffolds is estimated using a subset of the annotation metrics and viral sequences are used to identify circular templates (i.e., likely complete circular viruses). These quality analyses were determined to best reflect established completeness metrics for both bacteria and viruses (*61, 62*). Finally, VIBRANT compiles all results into a user-friendly format for visualization and downstream analysis. For a detailed description of VIBRANT’s workflow see Methods.

**Figure 1.**
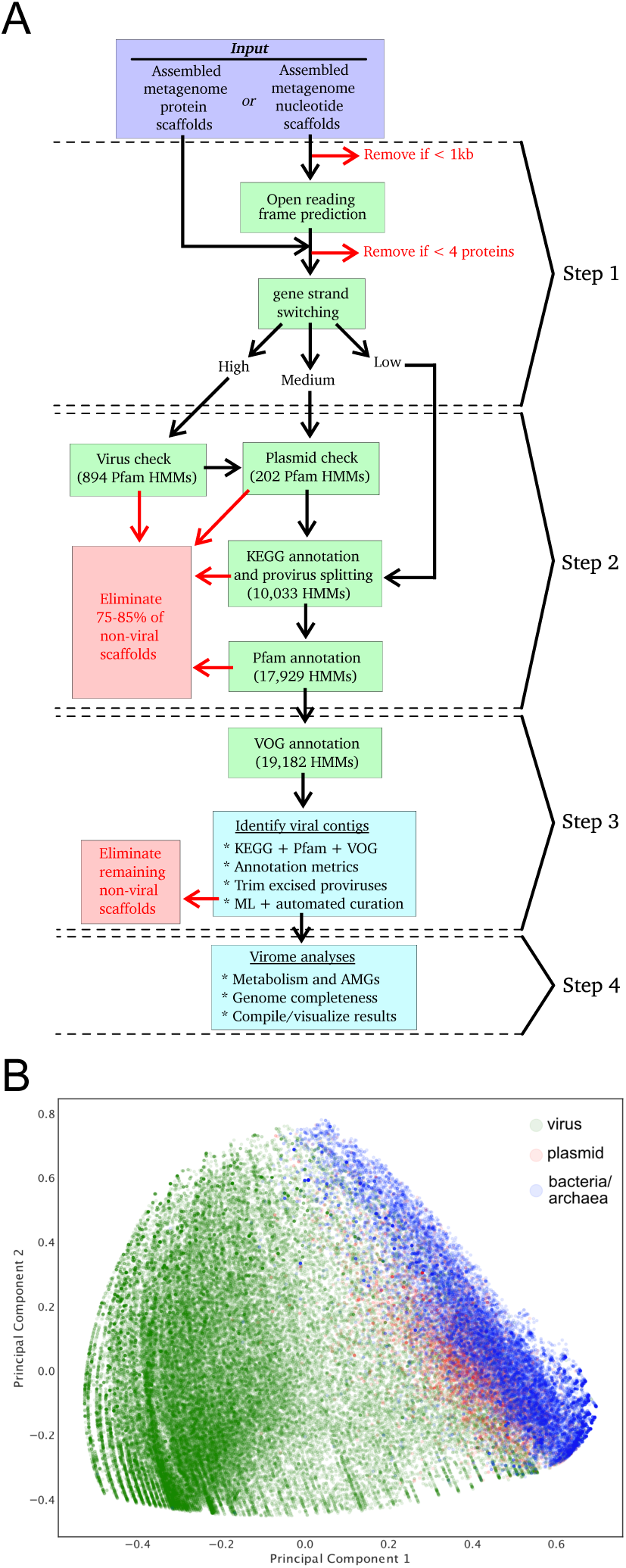
Representation of VIBRANT’s method for virus identification and virome functional characterization. (A) Workflow of virome analysis. Protein HMMs for analysis from KEGG, Pfam and VOG databases were used to construct signatures of viral and non-viral annotation metrics. (B) Visual representation (PCA plot) of the metrics used by the neural network to identify viruses, depicting viral, plasmid and bacterial/archaeal genomic sequences.

### Comparison of VIBRANT to other programs

VirSorter and VirFinder, two commonly used programs for identifying bacterial and archaeal viruses from metagenomes, were selected to compare against VIBRANT for the ability to accurately identify viruses. We evaluated all three programs’ performance on the same viral, bacterial and archaeal genomic, and plasmid datasets. Given that both VirSorter and VirFinder produce various confidence ranges of virus identification, we selected certain parameters for each program for comparison. For VirSorter, the parameters selected were [1] category 1 and 2 predictions, and [2] categories 1, 2 and 3 (i.e., all) predictions. For VirFinder, the intervals were [1] scores greater than or equal to 0.90 (approximately equivalent to a p-value of 0.013), and [2] scores greater than or equal to 0.75 (approximately equivalent to a p-value of 0.037). Hereafter, we provide two statistics for each VirSorter and VirFinder run that reflect results according to the two set confidence intervals, respectively.

VIBRANT yields a single output of confident predictions and therefore does not provide multiple output options. Since VIBRANT is only partially reliant on its neural network machine learning model for making predictions, all comparisons are focused on VIBRANT’s full workflow performance. VIBRANT does not consider scaffolds shorter than 1000 bp or those that encode less than four predicted open reading frames in order to maintain a low false positive rate (FPR) and have sufficient annotation information for identifying viruses. Therefore, in comparison of performance metrics only scaffolds meeting VIBRANT’s minimum requirements were analyzed. Inclusion of fragments encoding less than four open reading frames in analyses, which are frequently generated by metagenomic assemblies, are discussed below. We used the following calculations to compare performance: recall, precision, accuracy, specificity and F1 score (Figure 2).

**Figure 2.**
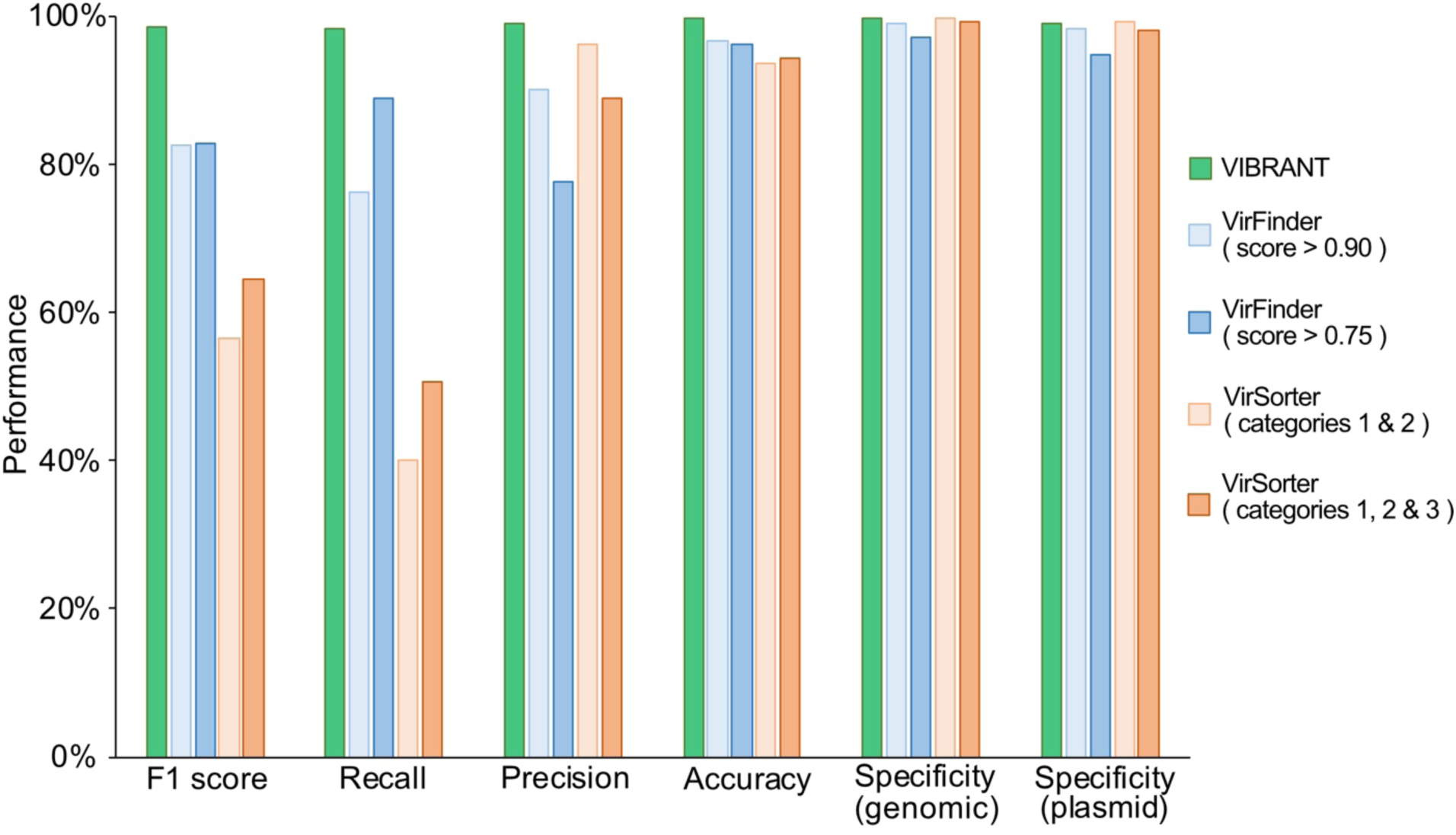
Performance comparison of VIBRANT, VirFinder and VirSorter on artificial scaffolds 3kb-15kb. Performance was evaluated using datasets of reference viruses, bacterial plasmids, and bacterial/archaeal genomes. For VirFinder and VirSorter two different confidence cutoffs were used (VirFinder: score of at least 0.90, and score of at least 0.75. VirSorter: categories 1 and 2 predictions, and categories 1, 2 and 3 predictions). All three programs were compared using the following statistical metrics: F1 score, recall, precision, accuracy and specificity. To ensure equal comparison all scaffolds tested encoded at least four open reading frames.

First, we evaluated the true positive rate (TPR, or recall) of viral genomic fragments as well as whole viral genomes. Viral genomes were acquired from the National Center for Biotechnology Information (NCBI) RefSeq and GenBank databases and split into various non-redundant fragments between 3 and 15 kb to simulate genomic scaffolds (Supplementary Table 1). VIBRANT correctly identified 98.38% of the 29,926 viral fragments, which was substantially greater than either VirSorter (40.00% and 50.67%) and VirFinder (76.23% and 89.02%).

Similar to TPR, we calculated FPR (or specificity) using two different datasets: genomic fragments of bacteria and archaea (hereafter genomic), and bacterial plasmids (plasmid). Plasmids were evaluated separately because they often encode for genes similar to those on viral genomes, such as those for genome replication and mobilization. Genomic and plasmid sequences were acquired from NCBI RefSeq and GenBank databases and split into various non-redundant fragments between 3 and 15 kb (Supplementary Table 1). Before analysis, putative proviruses were depleted from the datasets (see Methods). With the exception of VirSorter set at categories 1 and 2 for the plasmid dataset, VIBRANT had the highest specificity for both genomic (99.92%) and plasmid fragments (99.04%). VirSorter had similar specificity for both genomic (99.84% and 99.33%) and plasmid (99.33% and 98.10%) datasets, but only VirFinder set to a score cutoff of 0.90 was fully comparable (genomic: 99.10%, plasmid: 98.40%). At a score cutoff of 0.75, VirFinder was slightly less specific (genomic: 97.19%, plasmid: 94.92%). Although VirFinder (set to a score cutoff of 0.90) and VIBRANT had a similar overall specificity, VirFinder identified 11.8 times more bacterial/archaeal scaffolds as viruses (false discoveries) compared to VIBRANT (2,311 and 196, respectively).

We used the results from TPR of viral fragments and FPR of non-viral genomic or plasmid fragments to calculate precision, accuracy and F1 score. VIBRANT outperformed VirFinder and VirSorter at either score criteria in both precision (99.01%) and accuracy (99.74%). F1 is a metric (maximum value of 1) accounting for both TPR and FPR, and therefore acts as a comprehensive evaluation of overall performance. Our calculation of F1 indicates that VIBRANT (0.99) is able to better identify viruses while subsequently reducing false identifications compared to VirSorter (0.57 and 0.65) or VirFinder (0.83 and 0.83).

### Identification of viruses in diverse environments

We next tested VIBRANT’s ability to successfully identify viruses from a diversity of environments. Using 120,834 viruses from the Integrated Microbial Genomes and Viruses (IMG/VR v2.0) database (*63, 64*), in which the source environment of viruses is categorized, we identified that VIBRANT is more robust in identifying viruses from all tested environments compared to VirFinder and VirSorter (Figure 3A). Excluding air, in which there were only 62 representative viruses, VIBRANT averaged 94.5% recall, substantially greater than VirFinder (29.2% and 48.1%) and VirSorter (54.4% and 56.0%). These results suggest that in comparison to other software, VIBRANT has no evident database or environmental biases and is fully capable of identifying viruses from a broad range of source environments. We also used a dataset of 13,203 viruses from the Human Gut Virome database (*65*) for additional comparison. The vast majority of viruses (~96%) in this dataset were assumed to infect bacteria. Although recall was diminished compared to IMG/VR datasets, VIBRANT (78.7%) nevertheless outperformed both VirFinder (31.7% and 62.8%) and VirSorter (41.9% and 46.5%) on this dataset.

**Figure 3.**
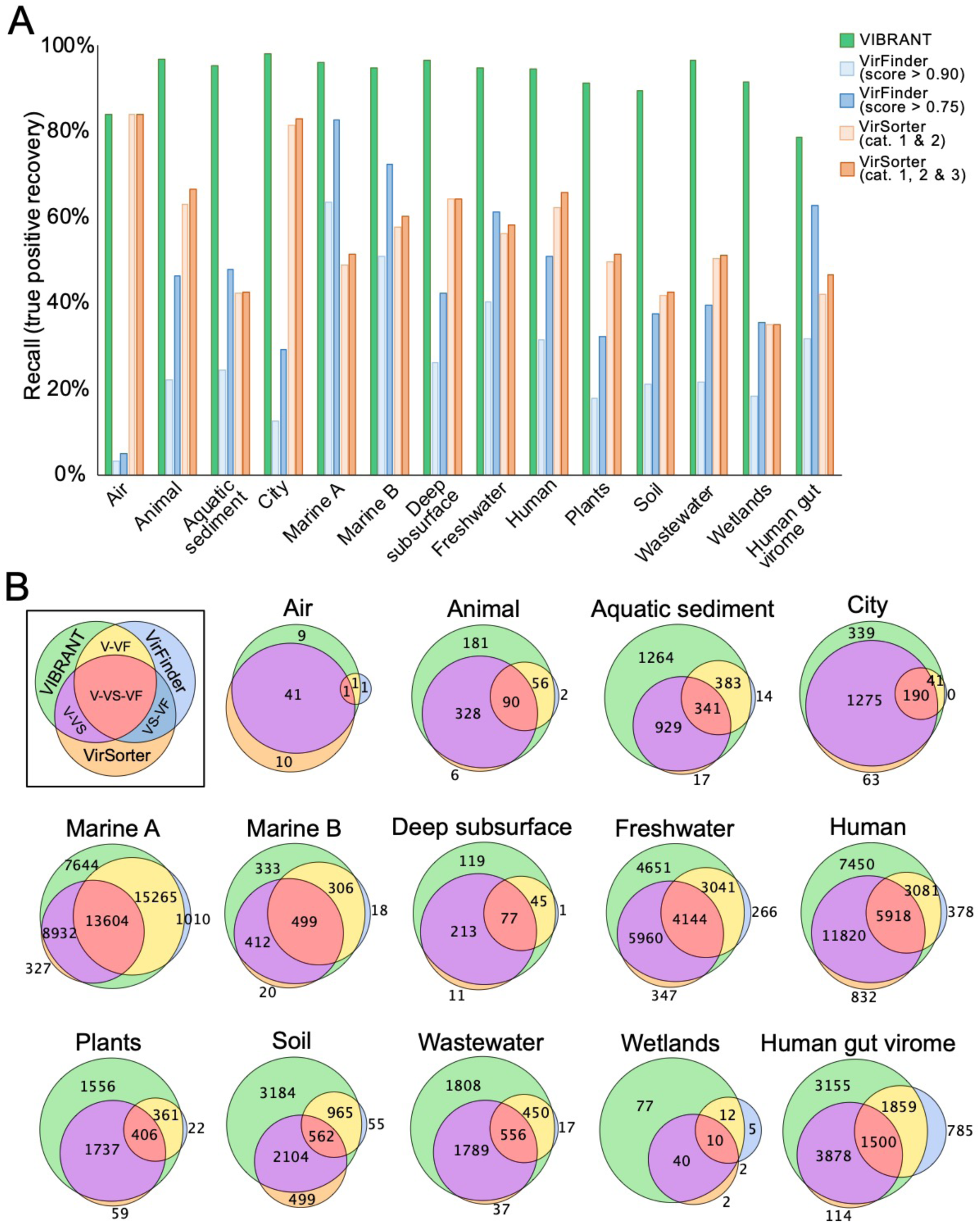
Effect of source environment on predictive abilities of VIBRANT, VirFinder and VirSorter. Viral scaffolds from IMG/VR and HGV database were used to test if VIBRANT displays biases associated with specific environments. (A) The recall (or recovery) of viral scaffolds from 14 environment groups was compared between VIBRANT and two confidence cutoffs for both VirFinder and VirSorter. Marine environments were classified into two groups: marine A (coastal, gulf, inlet, intertidal, neritic, oceanic, pelagic and strait) and marine B (hydrothermal vent, volcanic and oil). (B) Comparison of the overlap in the scaffolds identified as viruses by all three programs. Unique scaffolds identified by each program are in green (VIBRANT), orange (VirSorter) and light blue (VirFinder). The size of the circles represents the relative size of the group.

Many viruses from the IMG/VR dataset that were identified by VIBRANT were not identified by either VirFinder or VirSorter, indicating that VIBRANT has the propensity for discovery of novel viruses (Figure 3B). For most environments, the majority of viruses identified by VirFinder were already identified by either VIBRANT or VirSorter. The differences in the overlap of identified viruses was not too distinctive in environments for which many reference viruses are available, such as marine. For more understudied environments, such as plants or wastewater, VIBRANT displayed near-complete overlap with VirFinder and VirSorter predictions in conjunction with identifying over 40% more viruses.

### Identification of viruses in mixed metagenomes

Metagenomes assembled using short read technology contain many scaffolds that do not meet VIBRANT’s minimum length requirements and therefore are not considered during analysis. Despite this, VIBRANT’s predictions contain more annotation information and greater total viral sequence length than tools built to identify short sequences, such as scaffolds with less than four open reading frames. VIBRANT, VirFinder (score cutoff of 0.90) and VirSorter (categories 1 and 2) were used to identify viruses from human gut, freshwater lake and thermophilic compost metagenome sequences (Table 1). In addition, alternate program settings—VIBRANT “virome” mode, VirFinder score cutoff of 0.75 and VirSorter “virome decontamination” mode—were used to identify viruses from an estuary virome dataset. Each metagenomic assembly was limited to sequences of at least 1000bp but no minimum open reading frame limit was set. For these metagenomes, 31% to 40% of the scaffolds were of sufficient length (at least four open reading frames) to be analyzed by VIBRANT; for the estuary virome 62% were of sufficient length. In comparison, 100% of scaffolds from each dataset were long enough to be analyzed by VirFinder. The ability of VirFinder to make a prediction with each scaffold is considered the major strength of the tool.

**Table 1.** Virus recovery of VIBRANT, VirFinder and VirSorter from mixed metagenomes and a virome. Mixed community assembled metagenomes from the human gut, thermophilic compost and a freshwater lake, as well as an estuary virome, were used to compare virus prediction ability between the three programs. For each assembly the scaffolds were limited to a minimum length of 1000bp. Only a subset of each dataset contained scaffolds encoding at least four open reading frames. VIBRANT, VirFinder (score minimum of 0.90) and VirSorter (categories 1 and 2) were compared by total viral predictions, total combined length of predicted viruses, and total combined proteins of predicted viruses.

| Metagenome | sequences total (>1kb) | sequences 4+ ORFs | Metric | VIBRANT | VirFinder (score>0.90) | VIBRANT vs. VirFinder | VirSorter (cat. 1 & 2) | VIBRANT vs. VirSorter |
| --- | --- | --- | --- | --- | --- | --- | --- | --- |
| human gut: adenoma | 34,883 | 11,360 | total putative viruses | 505 | 604 | 0.84 | 284 | 1.78 |
|  |  |  | total virus length (bp) | 5,159,390 | 1,696,118 | 3.04 | 3,982,292 | 1.30 |
|  |  |  | total virus proteins | 7,534 | 2,134 | 3.53 | 5,484 | 1.37 |
| human gut: carcinoma | 53,946 | 18,669 | total putative viruses | 744 | 1,329 | 0.56 | 450 | 1.65 |
|  |  |  | total virus length (bp) | 5,415,994 | 3,500,838 | 1.55 | 4,182,862 | 1.29 |
|  |  |  | total virus proteins | 8,108 | 4,644 | 1.75 | 5,945 | 1.36 |
| human gut: healthy | 42,739 | 17,079 | total putative viruses | 548 | 672 | 0.82 | 309 | 1.77 |
|  |  |  | total virus length (bp) | 5,468,452 | 2,411,049 | 2.27 | 4,512,571 | 1.21 |
|  |  |  | total virus proteins | 7,998 | 3,230 | 2.48 | 6,127 | 1.31 |
| thermophilic compost | 68,815 | 21,620 | total putative viruses | 1,057 | 878 | 1.20 | 383 | 2.76 |
|  |  |  | total virus length (bp) | 6,577,000 | 2,238,129 | 2.94 | 3,290,654 | 2.00 |
|  |  |  | total virus proteins | 9,908 | 2,806 | 3.53 | 4,400 | 2.25 |
| freshwater lake (bog) | 79,862 | 26,832 | total putative viruses | 5,600 | 7,567 | 0.74 | 1,503 | 3.73 |
|  |  |  | total virus length (bp) | 34,861,470 | 25,357,664 | 1.37 | 15,436,797 | 2.26 |
|  |  |  | total virus proteins | 55,976 | 37,537 | 1.49 | 21,280 | 2.63 |
| * estuary virome | 5,247 | 3,277 | total putative viruses | 3,135 | 2,294 | 1.37 | 1,121 | 2.80 |
|  |  |  | total virus length (bp) | 10,241,625 | 6,478,804 | 1.58 | 5,163,674 | 1.98 |
|  |  |  | total virus proteins | 20,475 | 12,035 | 1.70 | 9,645 | 2.12 |

\* VIBRANT, VirFinder and VirSorter ran with alternate settings

For all six assemblies VirFinder averaged approximately 1.2 times more virus identifications than VIBRANT, though for both thermophilic compost and the estuary virome VIBRANT identified a greater number. Despite VirFinder averaging more total virus identifications, VIBRANT averaged just over 2.1 times more total viral sequence length and 2.4 times more total viral proteins. This is the result of VIBRANT having the capability to identify more viruses of higher quality and longer sequence length. For example, among all six datasets VIBRANT identified 1,309 total viruses at least 10 kb in length in comparison to VirFinder’s 479. VIBRANT was also able to outperform VirSorter in all metrics, averaging 2.4 times more virus identifications, nearly 1.7 times more total viral sequence length, and 1.8 times more encoded viral proteins.

VIBRANT’s method of predicting viral scaffolds provides a unique opportunity in comparison to similar tools in that it yields scaffolds of higher quality which are more amenable for analyzing protein function in viromes. It is an important distinction that the total number of viruses identified may not be correlated with the total viral sequence identified or the total number of encoded proteins. Even if VIBRANT identified fewer total viral sequences compared to other tools in certain circumstances, more data of higher quality was generated as viral sequences of longer length were identified as compared to many short fragments. This provides an important distinction that the metric of total viral predictions is not necessarily an accurate representation for the quality or quantity of the data generated.

### Integrated provirus prediction

In many environments, integrated proviruses can account for a substantial portion of the active viral community (*66*). Despite this, few tools exist that are capable of identifying both lytic viruses from metagenomic scaffolds as well as proviruses that are integrated into host genomes. To account for this important group of viruses, VIBRANT identifies provirus regions within metagenomic scaffolds or whole genomes. VIBRANT is unique from most provirus prediction tools in that it does not rely on sequence motifs, such as integration sites, and therefore is especially useful for partial metagenomic scaffolds in which neither the provirus nor host region is complete. In addition, this functionality of VIBRANT provides the ability to trim non-viral (i.e., host genome) ends from viral scaffolds. This results in a more correct interpretation of genes that are encoded by the virus and not those that are misidentified as being within the viral genome region. Briefly, VIBRANT identifies proviruses by first identifying and isolating scaffolds and genomes at regions spanning several annotations with low v-scores. These regions were found to be almost exclusive to host genomes. After cutting the original sequence at these regions, a refinement step trims the putative provirus fragment to the first instance of a virus-like annotation to remove leftover host sequence (Figure 4A). The final scaffold fragment is then analyzed by the neural network similar to non-excised scaffolds.

**Figure 4.**
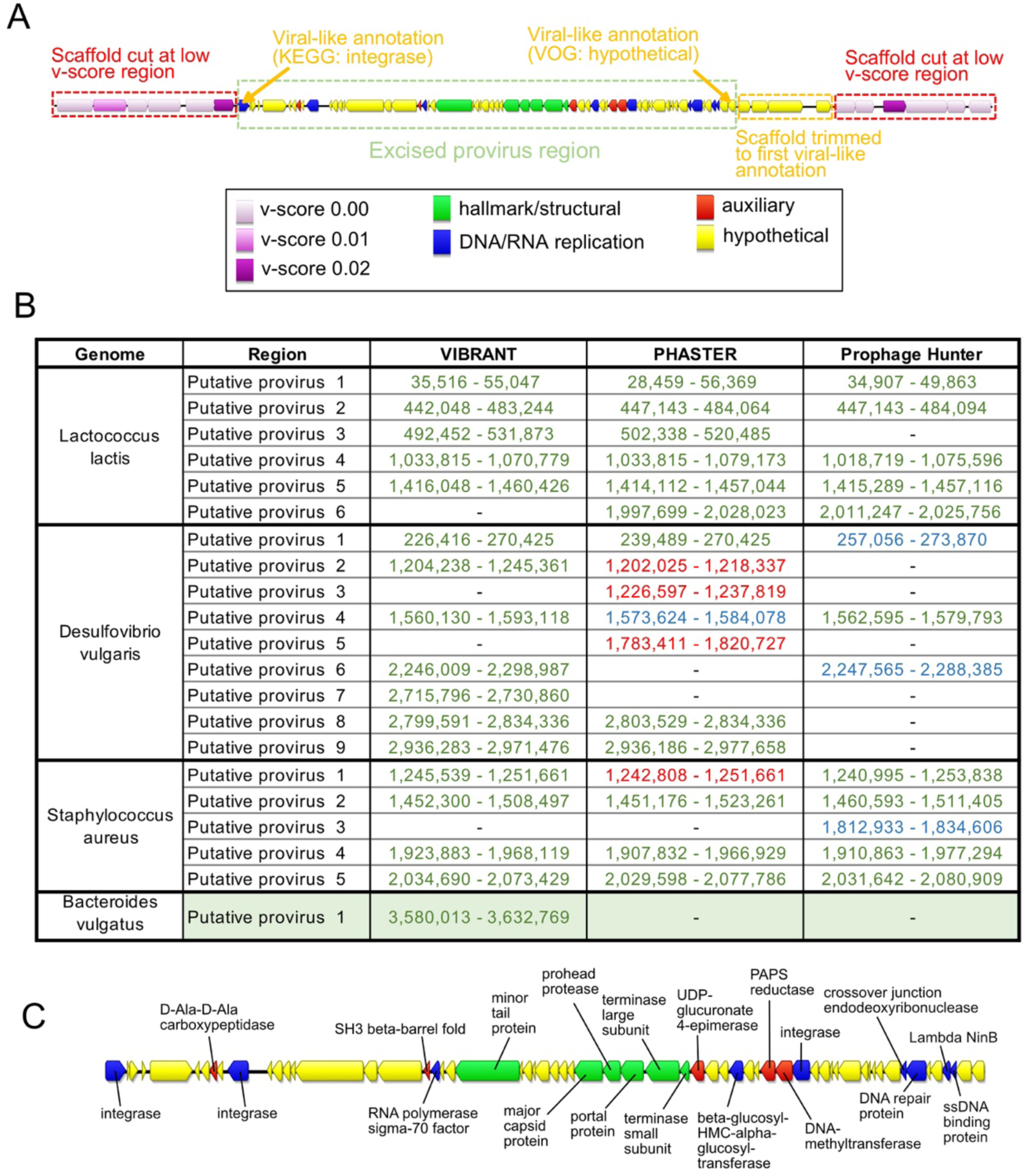
Prediction of integrated proviruses by VIBRANT, and comparison to PHASTER and Prophage Hunter. (A) Schematic representing the method used by VIBRANT to identify and extract provirus regions from host scaffolds using annotations. Briefly, v-scores are used to cut scaffolds at host-specific sites and fragments are trimmed to the nearest viral annotation. (B) Comparison of proviral predictions within four complete bacterial genomes between VIBRANT, PHASTER and Prophage Hunter. For PHASTER, putative proviruses are colored according to “incomplete” (red), “questionable” (blue) and “intact” (green) predictions. Prophage Hunter is colored according to “active” (green) and “ambiguous” (blue) predictions. (C) Manual validation of the *Bacteroides vulgatus* provirus prediction made by VIBRANT. The presence of viral hallmark protein, integrase and genome replication proteins strongly suggests this is an accurate prediction.

To assess VIBRANT’s ability to accurately extract provirus regions we compared its performance to PHASTER and Prophage Hunter, two programs explicitly built for this task. We compared the performance of these programs with VIBRANT on four bacterial genomes. VIBRANT and PHASTER predicted an equal number of proviruses, 17, while Prophage Hunter identified less, 13 (Figure 4B). Only one putative provirus prediction (*Lactococcus lactis* putative provirus 6) was shared between PHASTER and Prophage Hunter but not VIBRANT. However, VIBRANT was able to identify two putative provirus regions (*Desulfovibrio vulgaris* putative provirus 7 and *Bacteroides vulgatus* putative provirus 1) that neither PHASTER nor Prophage Hunter identified. Manual inspection of the putative *Bacteroides vulgatus* provirus identified a number of *bona fide* virus hallmark and virus-like proteins suggesting that it is an accurate prediction (Figure 4C). Our results suggest VIBRANT has the ability to accurately identify proviruses and, in some cases, can outperform other tools in this task.

### Evaluating quality of viral scaffolds and genomes

Determination of quality, in relation to completeness, of a viral scaffold has been notoriously difficult due to the absence of universally conserved viral genes. To date the most reliable metric of completeness for metagenomically assembled viruses is to identify circular sequences (i.e., complete circular genomes). Therefore, the remaining alternatives rely on estimation based on encoded proteins that function in central viral processes: replication of genomes and assembly of new viral particles.

VIBRANT estimates the quality of predicted viral scaffolds, a relative proxy for completeness, and indicates scaffolds that are circular. To do this, VIBRANT uses annotation metrics of nucleotide replication and viral hallmark proteins. Hallmark proteins are those typically specific to viruses and those that are required for productive infection, such as structural (e.g., capsid, tail, baseplate), terminase or viral holin/lysin proteins. Nucleotide replication proteins are a variety of proteins associated with either replication or metabolism, such as nucleases, polymerases and DNA/RNA binding proteins. Genomic scaffolds are categorized as low, medium or high quality draft as determined by VOG annotations (Figure 5A, Supplementary Table 2). High quality draft represents scaffolds that are likely to contain the majority of a virus’s complete genome and will contain annotations that are likely to aid in analysis of the virus, such as phylogenetic relationships and true positive verification. Medium draft quality represents the majority of a complete viral genome but is more likely to be a smaller portion in comparison to high quality. These scaffolds may contain annotations useful for analysis but are under less strict requirements compared to high quality. Finally, low draft quality constitutes scaffolds that were not found to be of high or medium quality. Many metagenomic scaffolds will likely be low quality genome fragments, but this quality category may still contain the higher quality genomes of some highly divergent viruses.

**Figure 5.**
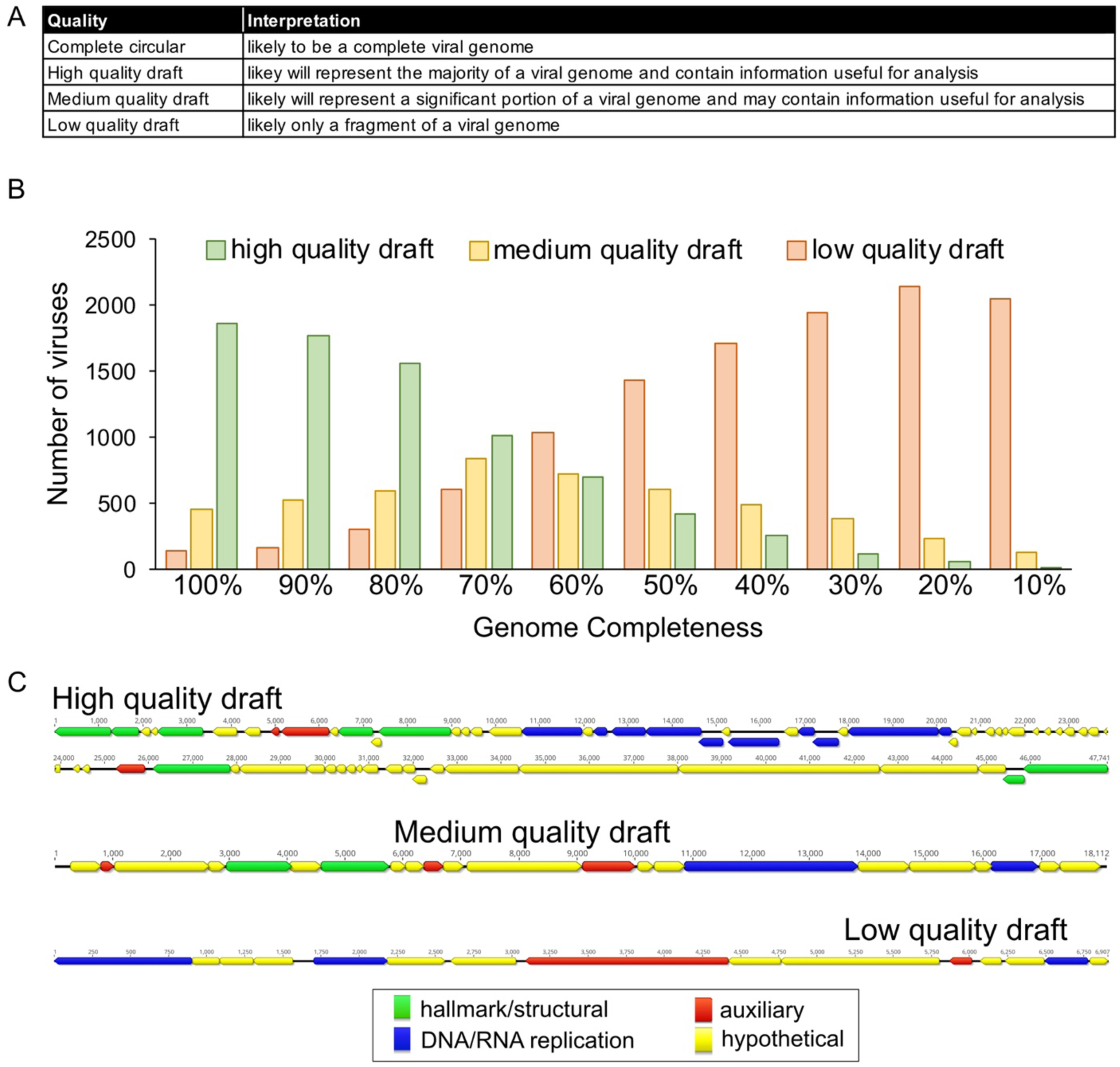
Estimation of genome quality of identified viral scaffolds. (A) Explanation of interpretation of quality categories: complete circular, high quality draft, medium quality draft and low quality draft. Quality generally represents total proteins, viral annotations, viral hallmark protein and nucleotide replication proteins, which are common metrics used for manual verification of viral genomes. (B) Application of quality metrics to 2466 NCBI RefSeq *Caudovirales* viruses with decreasing genome completeness from 100% to 10% completeness, respective of total sequence length. All 2466 viruses are represented within each completeness group. (C) Examples of viral scaffolds representing low, medium and high quality draft categories.

We benchmarked VIBRANT’s viral genome quality estimation using a total of 2466 *Caudovirales* genomes from NCBI RefSeq database. Genomes were evaluated either as complete sequences or by removing 10% of the sequence at a time stepwise between 100% and 10% completeness (Figure 5B). The results of VIBRANT’s quality analysis displayed a linear trend in indicating more complete genomes as high quality and less complete genomes as lower quality. The transition from categorizing genomes as high quality to medium quality ranged from 60% and 70% completeness. Although we acknowledge that VIBRANT’s metrics are not perfect, we demonstrate the first benchmarked approach to quantify and characterize genome quality associated with completeness of viral scaffolds. Manual inspection and visual verification of viral genomes that were characterized into each of these genome quality categories showed that quality estimations matched annotations (Figure 5C).

### Identifying function in virome: metabolic analysis

Viruses are a dynamic and key facet in the metabolic networks of microbial communities and can reprogram the landscape of host metabolism during infection. This can often be achieved by modulating host metabolic networks through expression of AMGs encoded on viral genomes. Identifying these AMGs and their associated role in the function of communities is imperative for understanding complex microbiome dynamics, or in some cases can be used to predict virus-host relationships. VIBRANT is optimized for the evaluation of function in viromes by identifying and classifying the metabolic capabilities of the viral community. To do this, VIBRANT identifies AMGs and assigns them into specific metabolic pathways and broader categories as designated by KEGG annotations.

To highlight the utility of this feature we compared the metabolic function of viruses derived from several diverse environments: freshwater, marine, soil, human-associated and city (Supplementary Figure 1). We found natural environments (freshwater, marine and soil) to display a different pattern of metabolic capabilities compared to human environments (human-associated and city). Viruses originating from natural environments tend to largely encode AMGs for amino acid and cofactor/vitamin metabolism with a more secondary focus on carbohydrate and glycan metabolism. On the other hand, AMGs from city and human environments are dominated by amino acid metabolism, and to some extent cofactor/vitamin and sulfur relay metabolism. In addition to this broad distinction, all five environments appear slightly different from each other. Despite freshwater and marine environments appearing similar in the ratio of AMGs by metabolic category, the overlap in specific AMGs is less extensive. The dissimilarity between natural and human environments is likewise corroborated by the relatively low overlap in individual AMGs.

A useful observation provided by VIBRANT’s metabolic analysis is that there appears to be globally conserved AMGs (i.e., present within at least 10 of the 13 environments tested). These 14 genes—*dcm, cysH, folE, phnP, ubiG, ubiE, waaF, moeB, ahbD, cobS, mec, queE, queD, queC*—likely perform functions that are central to viral replication regardless of host or environment. Notably, *folE*, *queD*, *queE* and *queC* constitute the entire 7-cyano-7-deazaguanine (preQ_0_) biosynthesis pathway, but the remainder of queuosine biosynthesis are entirely absent with the exception of *queF*. Certain AMGs are unique in that they are the only common representatives of a pathway amongst all AMGs identified, such as *phnP* for methylphosphonate degradation. These AMGs may indicate an evolutionary advantage for manipulating a specific step of a pathway, such as overcoming a reaction bottleneck, as opposed to modulating an entire pathway as seen with preQ_0_ biosynthesis. However, it should be noted that this list of 14 globally conserved AMGs may not be entirely inclusive of the core set of AMGs in a given environment.

VIBRANT was evaluated for its ability to provide new insights into virome function by highlighting AMGs from mixed metagenomes. Using only data from VIBRANT’s direct outputs, we compared the viral metabolic profiles of 6 hydrothermal vent and 15 human gut metagenomes (Figure 6). As anticipated, based on IMG/VR environment comparisons, the metabolic capabilities between the two environments were different even though the number of unique AMGs was relatively equal (138 for hydrothermal vents and 151 for human gut). The pattern displayed by metabolic categories for each metagenome was similar to that displayed by marine and human viromes. For hydrothermal vents the dominant AMGs were part of carbohydrate, amino acid and cofactor/vitamin metabolism, whereas human gut AMGs were mostly components of amino acid and, to some extent, cofactor/vitamin metabolism. Although the observed AMGs and metabolic pathways were overall different, about a third (50 total AMGs) of all AMGs from each environment were shared; between these metagenomes alone all 14 globally conserved AMGs were present.

**Figure 6.**
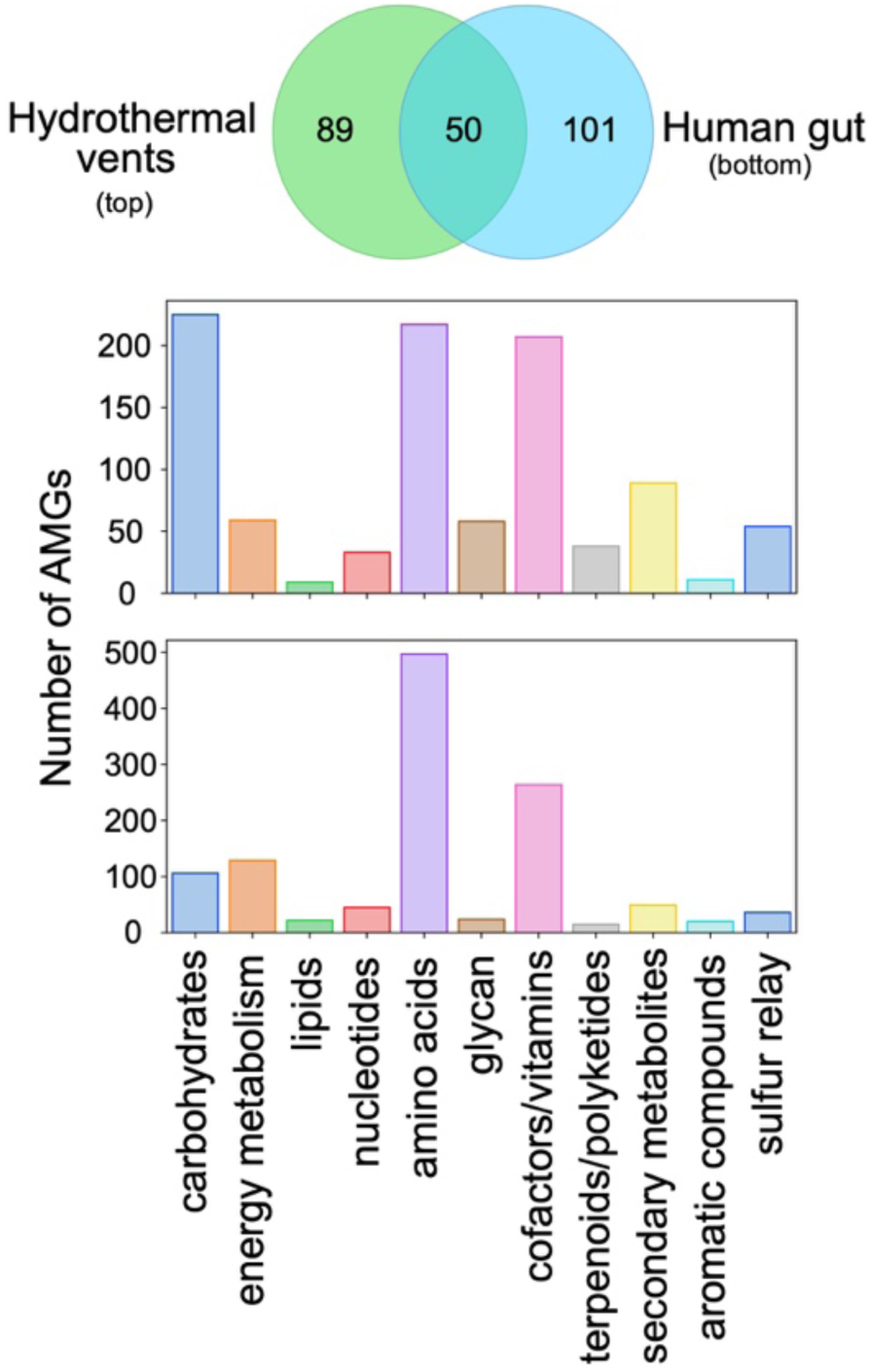
Comparison of AMG metabolic categories between hydrothermal vents and human gut. The Venn diagram depicts the unique and shared nonredundant AMGs between 6 hydrothermal vent and 15 human gut metagenomes. The graphs depict the differential abundance of KEGG metabolic categories of respective AMGs for hydrothermal vents (top) and human gut (bottom).

Observations of individual AMGs provided insights into how viruses interact within different environments. For example, tryptophan 7-halogenase (*prnA*) was identified in high abundance (45 total AMGs) within hydrothermal vent metagenomes but was absent from the human gut. Verification using GOV2 (Global Ocean Viromes 2.0) (*67*) and Human Gut Virome databases supported our finding that *prnA* appears to be constrained to aquatic environments, which is further supported by the gene’s presence on several marine cyanophages. PrnA catalyzes the initial reaction for the formation of pyrrolnitrin, a strong antifungal antibiotic. Identification of this AMG only within aquatic environments suggests a directed role in aquatic virus lifestyles. Similarly, cysteine desulfhydrase (*iscS*) was abundant (14 total AMGs) within the human gut metagenomes but not hydrothermal vents.

### Application of VIBRANT: Identification of viruses from individuals with Crohn’s Disease

We applied VIBRANT to identify viruses of at least 5kb in length from 102 human gut metagenomes (discovery dataset): 49 from individuals with Crohn’s Disease and 53 from healthy individuals (*68, 69*). VIBRANT identified 14,121 viruses out of 511,977 total scaffolds. These viral scaffolds were dereplicated to 8,822 non-redundant viral genomes using a cutoff of 95% nucleotide identity over at least 70% of the scaffold. We next used read coverage of each virus from all 102 metagenomes to calculate relative differential abundance across Crohn’s Disease and healthy individuals. In total, we found 721 viruses to be more abundant in the gut microbiomes associated with Crohn’s Disease (Crohn’s-associated) and 950 to be more abundant in healthy individuals (healthy-associated).

Using these viruses identified by VIBRANT we sought to identify taxonomic or host-association relationships to differentiate the virome of individuals with Crohn’s Disease. We used vConTACT2 to cluster the 721 Crohn’s- or 950 healthy-associated viruses with reference genomes using protein similarity. The majority of viruses (95.5%) were not clustered with any reference genome at approximately the genus level suggesting VIBRANT may have identified a large pool of novel or unique viral genomes. Although fewer total viruses were associated with Crohn’s Disease, significantly more were clustered to at least one representative at the genus level (72 for Crohn’s and 4 for healthy). Interestingly, no differentially abundant viruses from healthy individuals clustered with Enterobacterales-infecting reference viruses (enteroviruses), yet the majority (60/76) of Crohn’s-associated viruses were clustered with known enteroviruses, such as Lambda-and Shigella-related viruses. The remaining 16 viruses mainly clustered with *Caudovirales* infecting *Lactococcus*, *Clostridium*, *Riemerella*, *Klebsiella* and *Salmonella* species, though *Microviridae* and a likely complete crAssphage were also identified. A significant proportion of all Crohn’s-associated viruses (250/721), and the majority of genus-level clustered viruses (42/76), were found to be integrated sequences within a microbial genomic scaffold but were able to be identified due to VIBRANT’s ability to excise proviruses.

We also generated a protein sharing network containing all 721 Crohn’s and 950 healthy-associated viruses, which corresponded to taxonomic and host relatedness (Figure 7A). This protein network identified two different clustering patterns: [1] overlapping Crohn’s and healthy-associated viral populations clustered with Firmicutes-like viruses which may be indicative of a stable gut virome; [2] Crohn’s-associated viruses clustered with Enterobacterales-like and Fusobacterium-like viruses which may be indicative of a state of dysbiosis. The presence of a greater diversity and abundance of Enterobacterales and Fusobacteria has previously been linked to Crohn’s Disease (*70, 71*), and therefore the presence of viruses infecting these bacteria may provide similar information.

**Figure 7.**
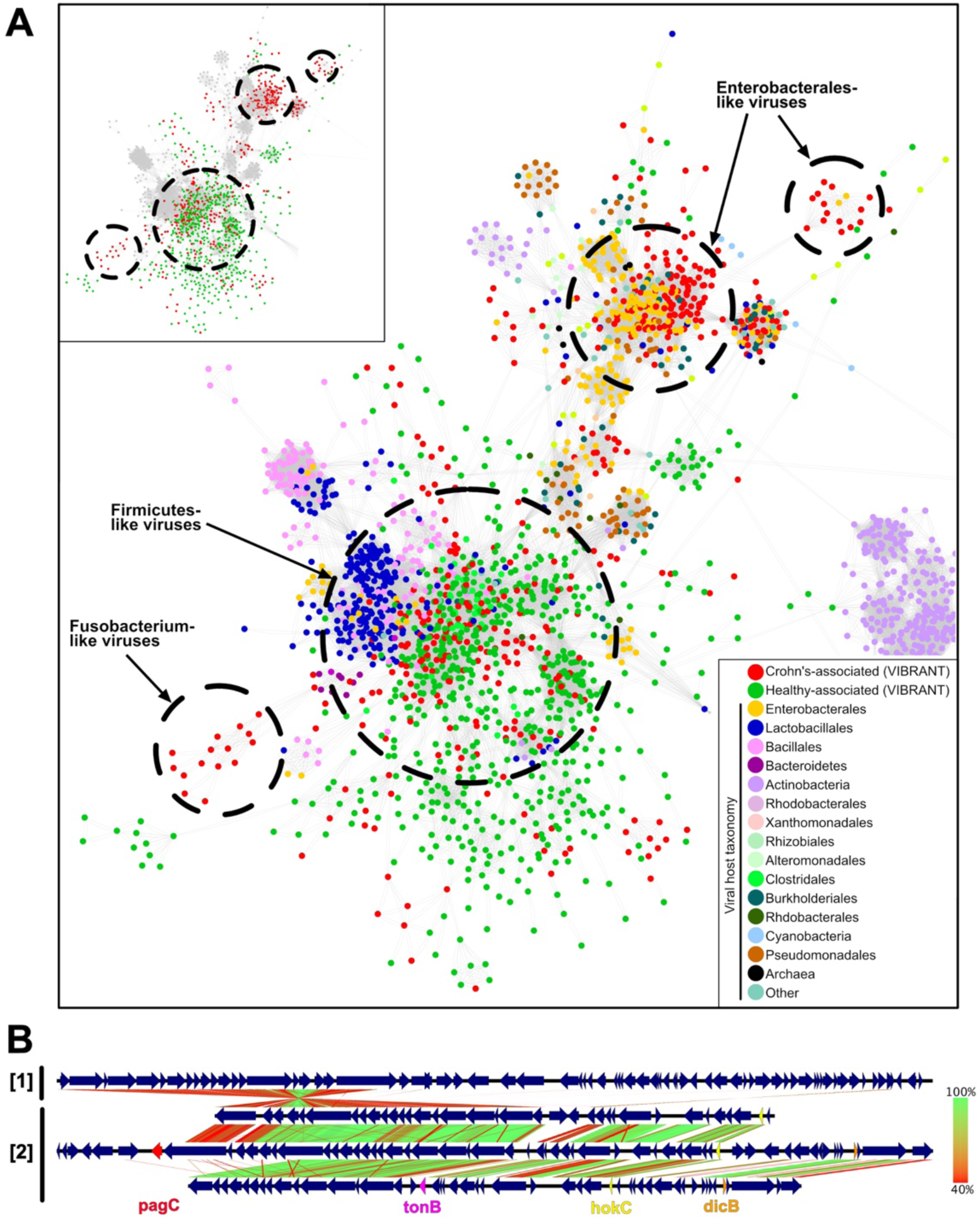
Viral metabolic comparison between Crohn’s Disease and healthy individuals gut metagenomes. (A) Partial view of vConTACT2 protein network clustering of viruses identified by VIBRANT and reference viruses. Small clusters and clusters with no VIBRANT representatives are not shown. Each dot represents one genome and is colored according to host or dataset association. Relevant viral groups are indicated by dotted circles (circles enclose estimated boundaries). (B) tBLASTx similarity comparison between [1] Escherichia phage Lambda and [2] three Crohn’s-associated viruses identified by VIBRANT. Putative virulence genes are indicated: *pagC*, *tonB*, *hokC* and *dicB*.

VIBRANT provides annotation information for all of the identified viruses which can be used to infer functional characteristics in conjunction with host association. Comparison of Crohn’s-associated Lambda-like virus genomic content and arrangement suggested a possible role of virally encoded host-persistence and virulence genes that are absent in the healthy-associated virome (Figure 7B). Among all Crohn’s-associated viruses, 17 total genes (*bor, dicB, dicC, hokC, kilR, pagC, ydaS, ydaT, yfdN, yfdP, yfdQ, yfdR, yfdS, yfdT, ymfL, ymfM* and *tonB*) that have the potential to impact host survival or virulence were identified. Importantly, no healthy-associated viruses encoded such genes (Table 2). The presence of these putative dysbiosis-associated genes (DAGs) may contribute to the manifestation and/or persistence of disease, similar to what has been proposed for the bacterial microbiome (*72–74*). For example, *pagC* encodes an outer membrane virulence factor associated with enhanced survival of the host bacterium within the gut (*75*). The identification of *dicB* encoded on a putative *Escherichia* virus is unique in that it may represent a ‘cryptic’ provirus that protects the host from lytic viral infection, thus likely to enhance the ability of the host to survive within the gut (*76*). Finally, *hokC* may indicate mechanisms of virally encoded virulence (*77*).

**Table 2.** Identification of putative DAGs encoded by Crohn’s-associated viruses. The differential abundance between Crohn’s Disease and healthy metagenomes of 17 putative DAGs. Abundance of each gene represents non-redundant annotations from Crohn’s-associated and healthy-associated viruses.

| ID | Gene | Name | Crohn's Disease | Healthy |
| --- | --- | --- | --- | --- |
| PF06291.11 | *bor* | Bor protein | 8 | 0 |
| K22304 | *dicB* | cell division inhibition protein | 8 | 0 |
| K22302 | *dicC* | transcriptional repressor of cell division inhibition gene *dicB* | 18 | 0 |
| K18919 | *hokC* | protein HokC/D | 16 | 0 |
| VOG11478 | *kilR* | Killing protein | 15 | 0 |
| K07804 | *pagC* | putative virulence related protein | 13 | 0 |
| PF15943.5 | *ydaS* | Putative antitoxin of bacterial toxin-antitoxin system | 22 | 0 |
| PF06254.11 | *ydaT* | Putative bacterial toxin | 18 | 0 |
| VOG04806 | *yfdN* | Uncharacterized protein | 19 | 0 |
| VOG01357 | *yfdP* | Uncharacterized protein | 11 | 0 |
| VOG11472 | *yfdQ* | Uncharacterized protein | 11 | 0 |
| VOG01639 | *yfdR* | Uncharacterized protein | 17 | 0 |
| VOG01103 | *yfdS* | Uncharacterized protein | 18 | 0 |
| VOG16442 | *yfdT* | Uncharacterized protein | 8 | 0 |
| VOG00672 | *ymfL* | Uncharacterized protein | 25 | 0 |
| VOG21507 | *ymfM* | Uncharacterized protein | 9 | 0 |
| K03832 | *tonB* | periplasmic protein | 3 | 0 |

To characterize the distribution and association of DAGs with Crohn’s Disease, we calculated differential abundance for two DAG-encoding viruses across all metagenome samples. The first virus encoded *pagC* and *yfdN*, and the second encoded *dicB*, *dicC* and *hokC*. Comparison of Crohn’s Disease to healthy metagenomes indicates these viruses are present within the gut metagenomes of multiple individuals but more abundant in association with Crohn’s Disease (Figure 8A). This suggests an association of disease with not only putative DAGs, but also specific, and potentially persistent, viral groups that encode them. In order to correlate increased abundance with biological activity we calculated the index of replication (iRep) for each of the two viruses (*78*). Briefly, iRep is a function of differential read coverage which is able to provide an estimate of active genome replication. Seven metagenomes containing the greatest abundance for each virus were selected for iRep analysis and indicated that each virus was likely active at the time of collection (Figure 8B).

**Figure 8.**
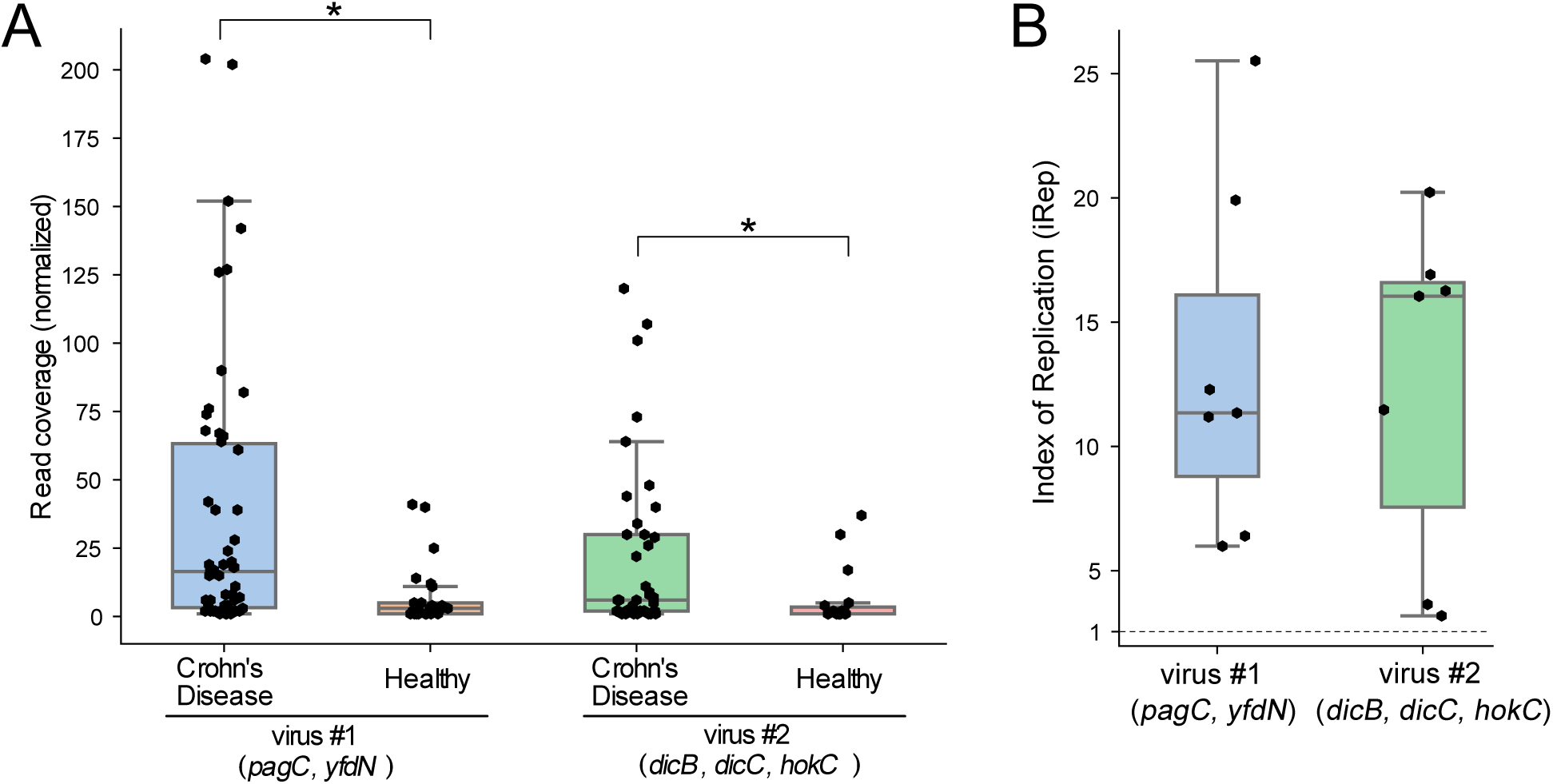
Differential abundance and activity of two viruses associated with Crohn’s Disease. (A) Normalized read coverage of two Crohn’s-associated viruses that encode putative DAGs between Crohn’s Disease and healthy gut metagenomes. Asterisks represent significant differential abundance (p<0.05). (B) iRep analysis for the same two viruses as (A), representative of seven metagenomes per virus. The dotted line indicates an iRep value of one, or low to no activity.

To validate these aforementioned findings, we applied VIBRANT to two additional metagenomic datasets from cohorts of individuals with Crohn’s disease and healthy individuals (validation dataset): 43 from individuals with Crohn’s Disease and 21 from healthy individuals (*79, 80*). VIBRANT identified 3,759 redundant viral genomes from Crohn’s-associated metagenomes and 1,444 from healthy-associated metagenomes. Determination of protein networks and visualization similarly identified clustering of Crohn’s-associated viruses with reference enteroviruses (Supplementary Figure 2). Likewise, we were able to identify 15 out of the 17 putative DAGs to be present in higher abundance in the Crohn’s Disease microbiome. This validates our findings of the presence of unique viruses and proteins associated with Crohn’s Disease, and suggests Enterobacterales-like viruses and putative DAGs may act as markers of Crohn’s Disease. Overall, our results suggest that VIBRANT provides a platform for characterizing these relationships.

## Discussion

Viruses that infect bacteria and archaea are key components in the structure, dynamics, and interactions of microbial communities. Tools that are capable of efficient recovery of these viral genomes from mixed metagenomic samples are likely to be fundamental to the growing applications of metagenomic sequencing and analyses. Importantly, such tools would need to reduce bias associated with specific viral groups (e.g., *Caudovirales*) and highly represented environments (e.g., marine). Moreover, viruses that exist as integrated proviruses within host genomes should not be ignored as they can represent a substantial fraction of infections in certain conditions and also persistent infections within a community.

Here we have presented VIBRANT, a novel method for the automated recovery of both free and integrated viral genomes from metagenomes that hybridizes neural network machine learning and protein signatures. VIBRANT utilizes metrics of non-reference based protein similarity annotation from KEGG, Pfam and VOG databases in conjunction with a novel ‘v-score’ metric to recover viruses with little to no biases. VIBRANT was built with the consideration of the human guided intuition used to manually inspect metagenomic scaffolds for viral genomes and packages these ideas into an automated software. This platform originates from the notion that proteins generally considered as non-viral, such as ribosomal proteins (*81*), may be decidedly common amongst viruses and should be considered accordingly when viewing annotations. V-scores are meant to provide a quantitative metric for the level of virus-association for each annotation used by VIBRANT, especially for Pfam and KEGG HMMs. That is, v-scores provide a means for both highlighting common or hallmark viral proteins as well as differentiating viral from non-viral annotations. In addition, v-scores give a quantifiable value to viral hallmark genes instead of categorizing them in a binary fashion.

VIBRANT was not only built for the recovery of viral genomes, but also to act as a platform for investigating the function of a virome. VIBRANT supports the analysis of virome function by assembling useful annotation data and categorizing the metabolic pathways of viral AMGs. Using annotation signatures, VIBRANT furthermore is capable of estimating genome quality and distinguishing between lytic and lysogenic viruses. To our knowledge, VIBRANT is the first software that integrates virus identification, annotation and estimation of genome completeness into a stand-alone program.

Benchmarking and validation of VIBRANT indicated improved performance compared to VirSorter and VirFinder, two commonly used programs for identifying viruses from metagenomes. This included a substantial increase in the relationship between true virus identifications (recall, true positive rate) and false non-virus identifications (specificity, false positive rate). That is, VIBRANT recovered more viruses with no discernable expense to false identifications. The result was that VIBRANT was able to recover an average of 2.4 and 1.7 more viral sequence from real metagenomes than VirFinder and VirSorter, respectively. When tested on metagenome-assembled viral genomes from IMG/VR representing diverse environments VIBRANT was found to have no perceivable environment bias towards identifying viruses. In comparison to provirus prediction tools, specifically PHASTER and Prophage Hunter, VIBRANT was shown to be proficient in identifying viral regions within bacterial genomes. This included the identification of a putative *Bacteroides* provirus that the other two programs were unable to identify. The importance of integrated provirus prediction was underscored in the analysis of Crohn’s Disease metagenomes since it was found that a significant proportion of disease related viruses were temperate viruses existing as host-integrated genomes.

VIBRANT’s method allows for the distinction between scaffold size and coding capacity in designating the minimum length of virus identifications. Traditionally, a cutoff of 5000 bp has been used to filter for scaffolds of a sufficient length for analysis. This is under the presumption that a longer sequence will be likely to encode more proteins. For example, this cutoff has been adopted by IMG/VR. However, we suggest a total protein cutoff of four open reading frames rather than sequence length cutoff to be more suitable for comprehensive characterization of the viral community. VIBRANT’s method works as a strict function of total encoded proteins and is completely agnostic to sequence length for analysis. Therefore, the boundary of minimum encoded proteins will support a more guided cutoff for quality control of virus identifications. For example, increasing the minimum sequence length to 5000 bp will have no effect on accuracy or ability to recall viruses since VIBRANT will only be considerate of the minimum total proteins, which is set to four. The result will be the loss of all 1000 bp to 4999 bp viruses that still encode at least four proteins. To visualize this distinction, we applied VIBRANT with various length cutoffs to the previously used estuary virome (see Table 1). Input sequences were stepwise limited from 1000 bp to 10000 bp (1000 bp steps) or four open reading frames to 13 open reading frames (one open reading frame steps) in length. Limiting to open reading frames indicated a reduced drop-off in total virus identifications and total viral sequence compared to a minimum sequence length limit (Supplementary Figure 3).

The output data generated by VIBRANT—protein/gene annotation information, protein/gene sequences, HMM scores and e-values, viral sequences in FASTA and GenBank format, indication of AMGs, genome quality, etc.—provides a platform for easily replicated pipeline analyses. Application of VIBRANT to characterize the function of Crohn’s-associated viruses emphasizes this utility. VIBRANT was not only able to identify a substantial number of viral genomes, but also provided meaningful information regarding putative DAGs, viral sequences for differential abundance calculation and genome alignment, viral proteins for clustering, and AMGs for metabolic comparisons.

## Conclusions

Our construction of the VIBRANT platform expands the current potential for virus identification and characterization from metagenomic and genomic sequences. When compared to two widely used software programs, VirFinder and VirSorter, we show that VIBRANT improves total viral sequence and protein recovery from diverse human and natural environments. As sequencing technologies improve and metagenomic datasets contain longer sequences VIBRANT will continue to outcompete programs built for short scaffolds (e.g., 500-3000 bp) by identifying more higher quality genomes. Our workflow, through the annotation of viral genomes, aids in the capacity to discover how viruses of bacteria and archaea may shape an environment, such as driving specific metabolism during infection or dysbiosis in the human gut. Furthermore, VIBRANT is the first virus identification software to incorporate annotation information into the curation of predictions, estimation of genome quality and infection mechanism (i.e., lytic vs lysogenic). We anticipate that the incorporation of VIBRANT into microbiome analyses will provide easy interpretation of viral data, enabled by VIBRANT’s comprehensive functional analysis platform and visualization of information.

## Methods

### Dataset for generation and comparison of metrics

To generate training and testing datasets sequences representing bacteria, archaea, plasmids and viruses were downloaded from NCBI databases (accessed July 2019) (Supplementary Table 3). For bacteria/archaea, 181 genomes from diverse phylogenetic groups were randomly chosen. Likewise, a total of 1,452 bacterial plasmids were chosen. For viruses, NCBI taxids associated with viruses that infect bacteria or archaea were used to download reference virus genomes, which were then limited to only sequences above 3kb. Sequences not associated with genomes, such as partial genomic regions, were manually removed. This resulted in 15,238 total viral genomes. All sequences were split into non-overlapping fragments between 3kb and 15kb to simulate metagenome assembled scaffolds (hereafter called *fragments*).

Integrated viruses are common in both bacteria and archaea. To address this for generating a dataset devoid of viruses, PHASTER (accessed July 2019) was used to predict putative integrated viruses in the 181 bacteria/archaea genomes. Using BLASTn (*82*), any fragments that had significant similarity (at least 95% identity, at least 3kb coverage and e-value < 1e-10) to the PHASTER predictions were removed as contaminant virus sequence. The new bacteria/archaea dataset was considered depleted of prophages, but not entirely devoid of contamination. Next, the datasets for bacteria/archaea and plasmids were annotated with KEGG, Pfam and VOG (hmmsearch (v3.1), e-value < 1e-5) (*83*) to further remove contaminant virus sequence. Plasmids were included because it was noted that the dataset appeared to contain virus sequences, possibly due to misclassification of episomal proviruses as plasmids. Using manual inspection of the KEGG, Pfam and VOG annotations any sequence that clearly belonged to a virus was removed. The final datasets consisted of 400,291 fragments for bacteria/archaea, 14,739 for plasmids, and 111,963 for viruses.

### V-score generation

Reference and database viral proteins were used to generate v-scores. To be consistent between all 15,238 viruses acquired from NCBI, proteins were predicted for all genomes using Prodigal (-p meta, v2.6.3) (*84*). All VOG proteins were added to this dataset, which resulted in a total of 633,194 proteins. Redundancy was removed from the generated viral protein dataset using cdhit (v4.6) (*85*) with a identify cutoff of 95%, which resulted in a total of 240,728 viral proteins (Supplementary Table 4). This was the final dataset used to generate v-scores. All KEGG HMM profiles to be used by VIBRANT (method described below) were used to annotate the viral proteins. A v-score for each KEGG HMM profile was determined by the number of significant (e-value < 1e-5) hits by hmmsearch, divided by 100, and a maximum value was set at 10 after division. The same v-score generation was done for Pfam and VOG databases. Any HMM profile with no significant hits to the virus dataset was given a v-score of zero. For KEGG and Pfam databases, any annotation that was given a v-score above zero and contained the keyword “phage” was given a minimum v-score of 1. To highlight viral hallmark genes, any annotation within all three databases with the keyword *portal, terminase, spike, capsid, sheath, tail, coat, virion, lysin, holin, base plate, lysozyme, head* or *structural* was given a minimum v-score of 1. Non-phage annotations (e.g., *phage shock protein*, *reovirus core-spike protein*) were not considered. The resulting v-scores are a metric of virus association (i.e., do not take into account virus specificity, or association with non-viruses) and are manually tuned to put greater weight on viral hallmark genes (Supplementary Table 5). Raw HMM table outputs can be found in Supplementary Tables 6, 7 and 8 for KEGG, Pfam and VOG, respectively.

### Databases used by VIBRANT

VIBRANT uses HMM profiles from three different databases: KEGG, Pfam and VOG (Supplementary Table 9). For Pfam all HMM profiles were used. To increase speed, KEGG and VOG HMM databases were reduced in size to contain only profiles likely to annotate the viruses of interest. For KEGG this was done by only retaining profiles considered to be relevant to “prokaryotes” as determined by KEGG. For VOG this was done by only retaining profiles that had at least one significant hit to an NCBI-acquired viral protein database using BLASTp. That is, any VOG HMM profile given a v-score of zero was removed. The resulting databases consisted of 10,033 HMM profiles for KEGG, 17,929 for Pfam, and 19,182 for VOG.

Two additional databases consisting of redundant Pfam HMM profiles were also generated. The first database consisted of virus annotations which were determined by a text search of “bacteriophage” to the Pfam database. Only HMM profiles with v-scores above zero were considered and those common to bacteria/archaea (e.g., glutaredoxin) were manually removed. This resulted in 894 virus specific HMMs. The second database consisted of common plasmid annotations. Proteins were predicted for the plasmid dataset using Prodigal (-p meta) and all Pfam HMMs with a v-score of zero were used to annotate the plasmid proteins (e-value < 1e-5). Any annotation with at least 50 hits was retained as a common plasmid HMM profile, which resulted in 202 common plasmid HMMs.

### Non-neural network steps and assembly of annotation metrics

VIBRANT utilizes several manually curated cutoffs in order to remove the bulk of non-virus input scaffolds before the neural network classifier is implemented. These steps will result in the assembly of 27 annotation metrics that are used by the neural network classifier for virus identification, which is followed by additional manually set cutoffs to curate the results.

First, open reading frames predicted by Prodigal (-p meta) or user input proteins are used to calculate the fraction of strand switching per scaffold (strand switches divided by total genes). Scaffolds are then classified as having either a *low* (5%), *medium* (5-35%) or *high* (>35%) level of strand switching. Scaffolds with a high level are annotated with the 894 virus-specific Pfam HMMs and only retained if there is at least one significant hit (score > 50). Throughout, scaffolds that are not retained are eliminated from further analysis. Scaffolds with a medium-level, and those with a high-level that passed the previous cutoff, are annotated with the 202 common plasmid Pfam HMMs and only retained if there are three or less significant hits (score > 50). Scaffolds with a low level are combined with those from high/medium that passed the previous cutoff(s).

Scaffolds are then annotated with the 10,033 KEGG-derived HMMs. Putative integrated provirus regions are extracted at this step by using sliding windows of either four or nine proteins at a time (step size = 1 protein). Within these windows scaffolds are fragmented according to v-scores and total KEGG annotations. Within the 4-protein window, scaffolds can be cut if [1] there are 0-1 unannotated proteins, 3-4 proteins with a v-score of 0-0.02 and a combined v-score of less than 0.06, or [2] three consecutive proteins with a v-score of 0 (considered as a 3-protein window). Scaffolds will also be cut using a 9-protein window if nine consecutive proteins are annotated. Finally, if the final two proteins on a scaffold each have a v-score of 0, the scaffold will be cut. Only scaffold fragments that contain at least 8 proteins are retained. Following provirus excision, several manual cutoffs are used to remove obvious non-viral scaffolds. Briefly, this is done by removing scaffolds with a high density of KEGG annotations (e.g., over 70% if less than 15 proteins or over 50% if greater than 15 proteins) or a high number of annotations with a v-score of 0 (e.g., over 15). V-scores are also used such that a scaffold that may be removed for having a high density of KEGG annotations will be retained if the v-score meets a specific threshold (e.g., average of 0.2).

Scaffolds that are retained are annotated by the 17,929 Pfam HMMs. In a similar manner to KEGG, scaffolds meeting set cutoffs for density and v-scores of Pfam HMMs are either retained or removed. For example, scaffolds with less than 15 total or density under 60% Pfam annotations are retained; a scaffold will be retained if it has greater than 60% Pfam annotations as well as an average v-score of at least 0.15. For both KEGG and Pfam cutoffs full details of every cutoff see Supplementary Table 10.

Following the aforementioned cutoff steps approximately 75-85% of non-viral scaffolds are removed. At this point scaffolds are annotated by the 19,182 VOG HMMs. Using VOG annotations and v-scores from KEGG and Pfam, putative proviruses that were cut during KEGG annotation are now trimmed to remove ends that may still contain host proteins. To do this, any scaffold previously cut is trimmed, at both ends, to either the first instance of a VOG annotation or the first v-score of at least 0.1.

Annotations from all three databases are used to assemble 27 metrics for the neural network classifier. Briefly the metrics are as follows: [1] total proteins, [2] total KEGG annotations, [3] sum of KEGG v-scores, [4] total Pfam annotations, [5] sum of Pfam v-scores, [6] total VOG annotations, [7] sum of VOG v-scores, [8] total KEGG integration related annotations (e.g., integrase), [9] total KEGG annotations with a v-score of zero, [10] total KEGG integration related annotations (e.g., integrase), [11] total Pfam annotations with a v-score of zero, [12] total VOG redoxin (e.g., glutaredoxin) related annotations, [13] total VOG non-integrase integration related annotations, [14] total VOG integrase annotations, [15] total VOG ribonucleotide reductase related annotations, [16] total VOG nucleotide replication (e.g., DNA polymerase) related annotations, [17] total KEGG nuclease (e.g., restriction endonuclease) related annotations, [18] total KEGG toxin/anti-toxin related annotations, [19] total VOG hallmark protein (e.g., capsid) annotations, [20] total proteins annotated by KEGG, Pfam and VOG, [21] total proteins annotated by Pfam and VOG only, [22] total proteins annotated by Pfam and KEGG only, [23] total proteins annotated by KEGG and VOG only, [24] total proteins annotated by KEGG only, [25] total proteins annotated by Pfam only, [26] total proteins annotated by VOG only, and [27] total unannotated proteins. Non-annotation features such as gene density, average gene length and strand switching were not used because they were found to decrease performance of the neural network classifier despite being differentiating features between bacteria/archaea and viruses; viruses tend to have shorter genes, less intergenic space and strand switch less frequently. This decreased performance is likely due to several reasons, such as errors associated with protein prediction (e.g., missed open reading frame leading to a large “intergenic” gap) or that scaffolds, due to being fragmented genomes in most cases, behave differently than the genome as a whole. For example, genomic regions encoding for large structural proteins will have a higher average gene size or a small window of virus proteins may have a greater average strand switching level compared to the whole genome.

### Training and testing VIBRANT

The bacteria/archaea genomic, plasmid and virus datasets described above were used to train and test the machine learning model. Scikit Learn libraries were used to assess various machine learning strategies to identify the best performing algorithm. Among support vector machines, neural networks and random forests, we found that neural networks lead to the most accurate and comprehensive identification of viruses. Therefore, Scikit Learn’s (*86*) supervised neural network multi-layer perceptron classifier (hereafter neural network) was used. The portion of VIBRANT up until the neural network classifier (i.e., KEGG, Pfam and VOG annotation) was used to compile the 27 annotation metrics for each of the three datasets. To account for differences in scaffold sizes all metrics were normalized (i.e., divided by) to the total number of proteins encoded by the scaffold. The first metric, for total proteins, was normalized to log base 10 of itself. Each metric was weighted equally, though it is worth noting that the removal of several metrics, mainly metrics 8-18, did not significantly impact the accuracy of model’s prediction. The normalized results were randomized and non-redundant portions of these results were taken for training or testing the neural network. It is important to note that the testing set here was not used as the comprehensive testing set for the entire workflow. In total, 93,913 fragments were used for training and 9,000 were used for testing the neural network (Supplementary Tables 11 and 12).

To comprehensively test the performance of VIBRANT in its entirety a new testing dataset was generated consisting of fragments from the neural network testing set as well as additional fragments non-redundant to the previous training dataset. This new testing dataset was comprised of 256,713 fragments from bacteria/archaea, 29,926 from viruses and 8,968 from plasmids. Each met the minimum size requirement of VIBRANT: at least four open reading frames. For comparison to VirFinder (v1.1) and VirSorter (v1.0.3), the latter testing dataset was used. Two intervals for VirFinder and VirSorter were used for comparison. For VirSorter, the intervals selected were [1] category 1 and 2 predictions, and [2] categories 1, 2 and 3 (i.e., all) predictions. VirSorter was ran using the “Virome” database. For VirFinder, the intervals were [1] scores greater than or equal to 0.90 (approximately equivalent to a p-value of 0.013), and [2] scores greater than or equal to 0.75 (approximately equivalent to a p-value of 0.037). All equations used can be found in Supplementary Table 13 and results used for the generation of Figure 1 can be found in Supplementary Table 14.

### AMG identification

KEGG annotations were used to classify potential AMGs (Supplementary Table 15). KEGG annotations falling under the “metabolic pathways” category as well as “sulfur relay system” were considered. Manual inspection was used to remove non-AMG annotations, such as *nrdAB* and *thyAX*. Other annotations not considered dealt with direct nucleotide to nucleotide conversions. All AMGs were associated with a KEGG metabolic pathway map.

### Completeness estimation

Scaffold completeness is determined based on four metrics: circularization of scaffold sequence, VOG annotations, total VOG nucleotide replication proteins and total VOG viral hallmark proteins (Supplementary Table 16). In order to be considered a complete genome a sequence must be identified as likely circular. A kmer-based approach is used to do this. Specifically, the first 20 nucleotides are compared to 20-mer sliding windows within the last 900bp of the sequence. If a complete match is identified the sequence is considered a circular template. Scaffolds can also be considered a low, medium or high quality draft. To benchmark completeness, NCBI RefSeq viruses identified as *Caudovirales*, limited to 10 kb in length, were used to estimate completeness by stepwise removing 10% viral sequence at a time (Supplementary Table 2). Viral genome diagrams to depict genome quality and completeness, as well as provirus predictions, were made using Geneious Prime 2019.0.3.

### Additional viral datasets and metagenomes

IMG/VR v2.0 (accessed July 2019) was downloaded and scaffolds originating from air, animal, aquatic sediment, city, marine A (coastal, gulf, inlet, intertidal, neritic, oceanic, pelagic and strait), marine B (hydrothermal vent, volcanic and oil), deep subsurface, freshwater, human, plants, soil wastewater and wetland environments were selected for analysis. Venn diagram visualization of virus predictions with this dataset was made using Matplotlib (v3.0.0) (*87*). Several published, assembled metagenomes from IMG/VR representing diverse environments were selected for comparing VIBRANT, Virsorter and VirFinder (IMG taxon IDs: 3300005281, 3300017813 and 3300000439). Fifteen publicly available datasets from the human gut were assembled for assessing VIBRANT and comparing the three programs (*88*). Reads can be found under NCBI BioProject PRJEB7774 (ERR688591, ERR688590, ERR688509, ERR608507, ERR608506, ERR688584, ERR688587, ERR688519, ERR688512, ERR688508, ERR688634, ERR688618, ERR688515, ERR688513, ERR688505). Reads were trimmed using Sickle (v1.33) (*89*) and assembled using metaSPAdes (v3.12.0 65) (*90*) (--meta -k 21,33,55,77,99). For hydrothermal vents, six publicly available hydrothermal plume samples were derived from Guaymas Basin (one sample) and Eastern Lau Spreading Center (five samples). Reads can be found under NCBI BioProject PRJNA314399 (SRR3577362) and PRJNA234377 (SRR1217367, SRR1217459, SRR1217564, SRR1217566, SRR1217452, SRR1217567, SRR1217465, SRR1217462, SRR1217460, SRR1217463, SRR1217565). Reads were trimmed using Sickle and assembled using metaSPAdes (--meta -k 21,33,55,77,99). Details of assembly and processing are outlined in Zhou *et al*. (*91*). For analysis of Crohn’s Disease metagenomes by VIBRANT, publicly available metagenomes were used; the metagenomes were sequenced by He *et al*., Ijaz *et al*. and Gevers *et al*., and assembled by Pasolli *et al*. (Supplementary Tables 17, 18 and 19).

### Analysis of Crohn’s Disease metagenomes

Metagenomic reads from He *et al.* were assembled by Pasolli *et al.* and used for analysis. VIBRANT (-l 5000) was used to predict viruses from 49 metagenomes originating from individuals with Crohn’s Disease and 53 from healthy individuals (102 total samples). A total of 14,121 viruses were identified. Viral scaffolds were dereplicated using Mash (*92*) and Nucmer (*93*) to 95% nucleotide identity and 70% scaffold coverage. The longest scaffold was kept as the representative for a total of 8,822 dereplicated viral scaffolds. A total of 96 read sets were used (59 Crohn’s Disease and 37 healthy), trimmed using Sickle and aligned to the dereplicated scaffolds using Bowtie2 (-N 1, v2.3.4.1) (*94*) and the resulting coverages were normalized to total reads. The normalized relative coverage of each scaffold for all 96 samples were compared using DESeq2 (*95*) (Supplementary Table 20). Scaffolds in significantly different abundance between Crohn’s Disease and control samples were determined by a p-value cutoff of 0.05. iRep (default parameters) (*78*) was used to estimate replication activity of two Crohn’s-associated viruses. EasyFig (v2.2.2) (*96*) was used to generate genome alignments of Escherichia phage Lambda (NCBI accession number NC_001416.1) and three Crohn’s-associated viruses. vConTACT2 (v0.9.8) was ran using default parameters on the CyVerse Discovery Environment platform. Putative hosts of Crohn’s-associated and healthy-associated was estimated using proximity of vConTACT2 protein clustering and BLASTp identity (NCBI non-redundant protein database, assessed October 2019). Two additional read sets from Gevers *et al.* (*80*) and Ijaz *et al.* (*79*) were likewise assembled by Pasolli *et al.*. VIBRANT (-l 5000 -o 10) was used to predict viruses from 43 metagenomes originating from individuals with Crohn’s Disease and 21 from healthy individuals (64 total samples). In contrast to the discovery dataset viral genomes were not dereplicated and differential abundance was not determined. Instead viruses from each group were directly clustered using vConTACT2. Abundances of DAGs in the validation set were normalized to total viruses. Protein networks were visualized using Cytoscape (v3.7.2) (*98*).

## Availability of data and materials

VIBRANT is implemented in Python and all scripts and associated files are freely available at https://github.com/AnantharamanLab/VIBRANT/. All data and genomic sequences used for analyses are publicly available; see Supplementary Tables 3, 17, 18 and 19 for study and accession names. Full protein networks generated by vConTACT2 for Crohn’s- and healthy-associated viruses are available in Supplementary Data 1 and 2, respectively. VIBRANT is also freely available for use as an application through the CyVerse Discovery Environment. To use the application visit https://de.cyverse.org/de/?type=apps&app-id=c2864d3c-fd03-11e9-9cf4-008cfa5ae621&system-id=de, and for more details see https://wiki.cyverse.org/wiki/display/DEapps/VIBRANT-1.0.1. Additional details of relevant data are available from the corresponding author on request.

## Acknowledgements

We thank Upendra Devisetty for his assistance with dockerizing and integrating VIBRANT as a web-based application in the CyVerse Discovery Environment.

## Contributors

K.K and K.A designed the study, performed all analyses and interpretation of data, and wrote the manuscript. Z.Z contributed to conceptualization of study design and reviewed the manuscript. All authors have reviewed and approved the final manuscript.

## Funding

We thank the University of Wisconsin - Office of the Vice Chancellor for Research and Graduate Education, University of Wisconsin – Department of Bacteriology, and University of Wisconsin – College of Agriculture and Life Sciences for their support.

## Conflicts of interest

The authors declare no competing interests.

## Supplementary Information

Supplementary Figures 1-3, Supplementary Tables 1-20 and Supplementary Data 1-2.

## List of Supplementary Figures

**Supplementary Figure 1. AMG and metabolic pathways between diverse environments.** VIBRANT was used to predict viruses from IMG/VR datasets and the identified metabolic pathways and AMGs were compared for freshwater, marine, soil, city and human-associated environments (graphs). The respective AMGs and their abundances were likewise compared (venn diagram).

**Supplementary Figure 2. Protein network of two Crohn’s Disease validation datasets.** VIBRANT was used to predict viruses from two datasets for validation of marker virus and putative DAG discovery. The resulting viruses were used to construct a protein network indicating Crohn’s-associated viruses clustering with enteroviruses more often than healthy-associated viruses.

**Supplementary Figure 3. Comparison of limiting to sequence length or open reading frames.** VIBRANT was used to predict viruses from an estuary virome and set to limit to either scaffold length or total encoded open reading frames. The (A) total virus identifications and (B) total viral sequence length were compared to show that limiting to open reading frames will typically yield more data.

## List of Supplementary Tables

**Supplementary Table 1. Number and sizes of sequence fragments used to train and test VIBRANT for viruses, plasmids, and bacteria and archaea.**

**Supplementary Table 2. Number of *Caudovirales* genomes and genomic fragments identified per quality estimation category, exact rules used to estimate genome quality and the interpretation of quality estimations.**

**Supplementary Table 3. List of NCBI accession numbers for bacterial and archaeal genomes, plasmids, and viral genomes used in this study.**

**Supplementary Table 4. Protein prediction coordinates of dereplicated proteins from NCBI viruses used to generate v-scores.**

**Supplementary Table 5. List of all KEGG, Pfam and VOG annotation names and associated v-scores (if greater than zero).**

**Supplementary Table 6. Unparsed HMM table output from KEGG annotations used to generate KEGG v-scores.**

**Supplementary Table 7. Unparsed HMM table output from Pfam annotations used to generate Pfam v-scores.**

**Supplementary Table 8. Unparsed HMM table output from VOG annotations used to generate VOG v-scores.**

**Supplementary Table 9. List of all HMM names used by VIBRANT.**

**Supplementary Table 10. Description of set cutoffs implemented before neural network machine learning analysis for KEGG and Pfam annotations.**

**Supplementary Table 11. Normalized data used to train the neural network machine learning classifier.**

**Supplementary Table 12. Normalized data used to test the neural network machine learning classifier.**

**Supplementary Table 13. Equations used for benchmarking analyses.**

**Supplementary Table 14. Calculations and results of benchmarking analyses.**

**Supplementary Table 15. List of all KEGG annotations determined as AMGs.**

**Supplementary Table 16. List of all VOG annotations determined as nucleotide replication-associated or viral hallmark-associated, which are used during prediction and quality estimation.**

**Supplementary Table 17. List of datasets used from He *et al.***

**Supplementary Table 18. List of datasets used from Ijaz *et al.***

**Supplementary Table 19. List of datasets used from Gevers *et al*.**

**Supplementary Table 20. Results from DESeq2 analysis for 8,789 non-redundant viruses from the Crohn’s Disease discovery dataset.**

## List of Supplementary Data

**Supplementary Data 1. Protein network generated by vConTACT2 for Crohn’s Disease discovery dataset.**

**Supplementary Data 2. Protein network generated by vConTACT2 for Crohn’s Disease validation dataset.**

## Notes

https://github.com/AnantharamanLab/VIBRANT

