## Supplementary Figure 1 for "VIBRANT: Automated recovery, annotation and curation of microbial viruses, and evaluation of virome function from genomic sequences"

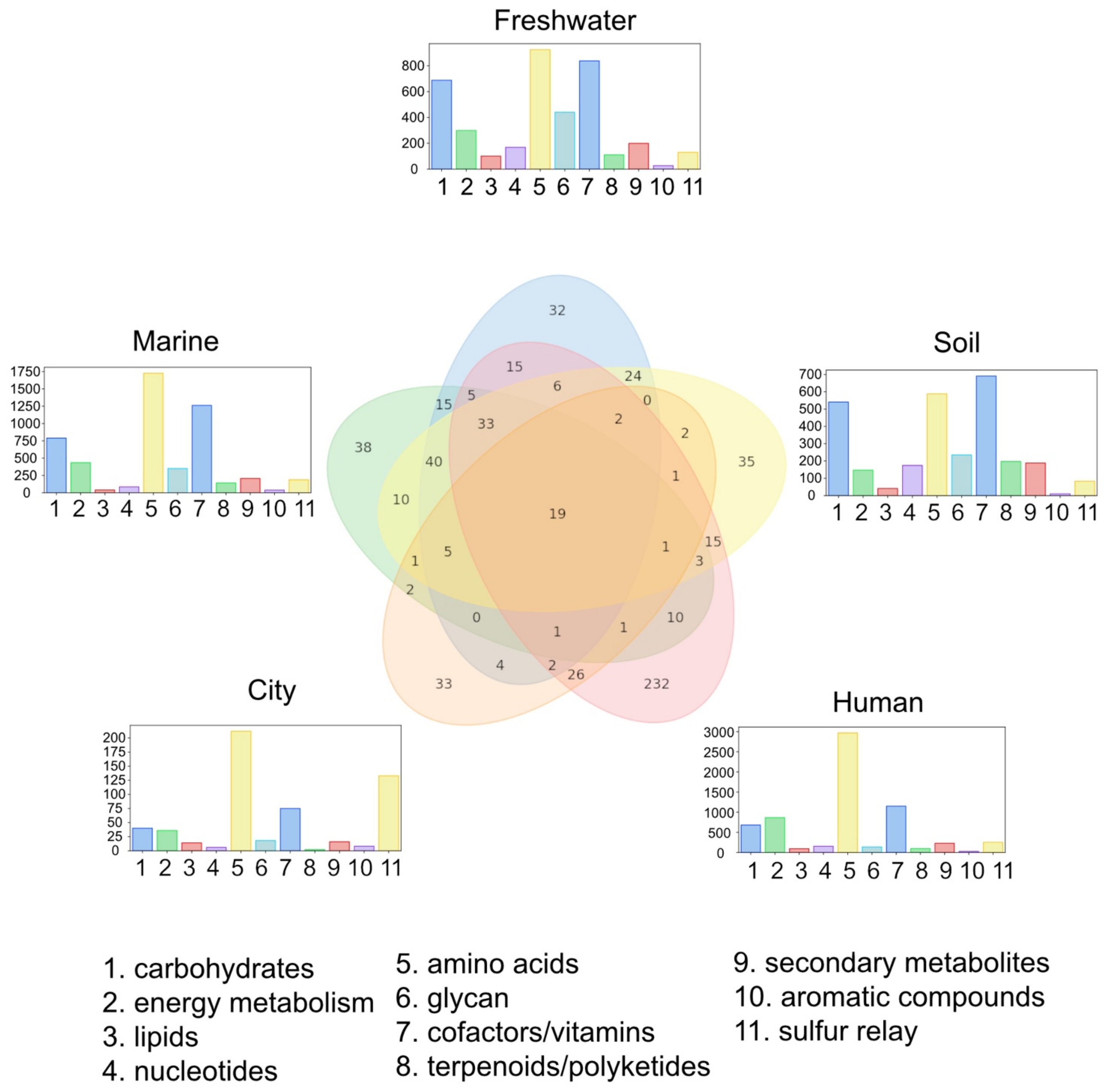


**Supplementary Figure 1. AMG and metabolic pathways between diverse environments.** VIBRANT was used to predict viruses from IMG/VR datasets and the identified metabolic pathways and AMGs were compared for freshwater, marine, soil, city and human-associated environments (graphs). The respective AMGs and their abundances were likewise compared (venn diagram).
