## Supplementary Figure 2 for "VIBRANT: Automated recovery, annotation and curation of microbial viruses, and evaluation of virome function from genomic sequences"

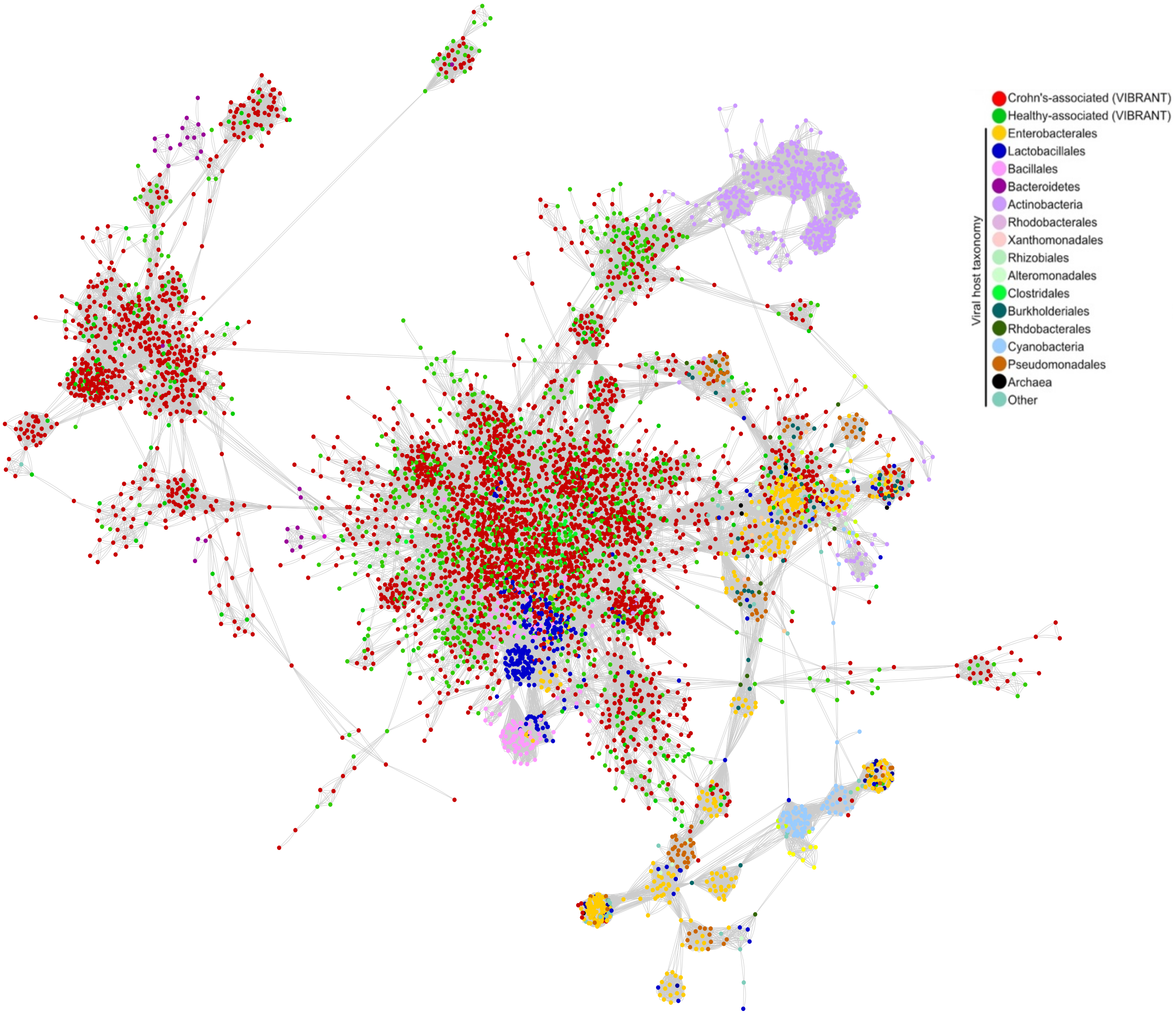


**Supplementary Figure 2. Protein network of two Crohn's Disease validation datasets.** VIBRANT was used to predict viruses from two datasets for validation of marker virus and putative DAG discovery. The resulting viruses were used to construct a protein network indicating Crohn's-associated viruses clustering with enteroviruses more often than healthy-associated viruses.
