## Supplementary Figure 3 for "VIBRANT: Automated recovery, annotation and curation of microbial viruses, and evaluation of virome function from genomic sequences"

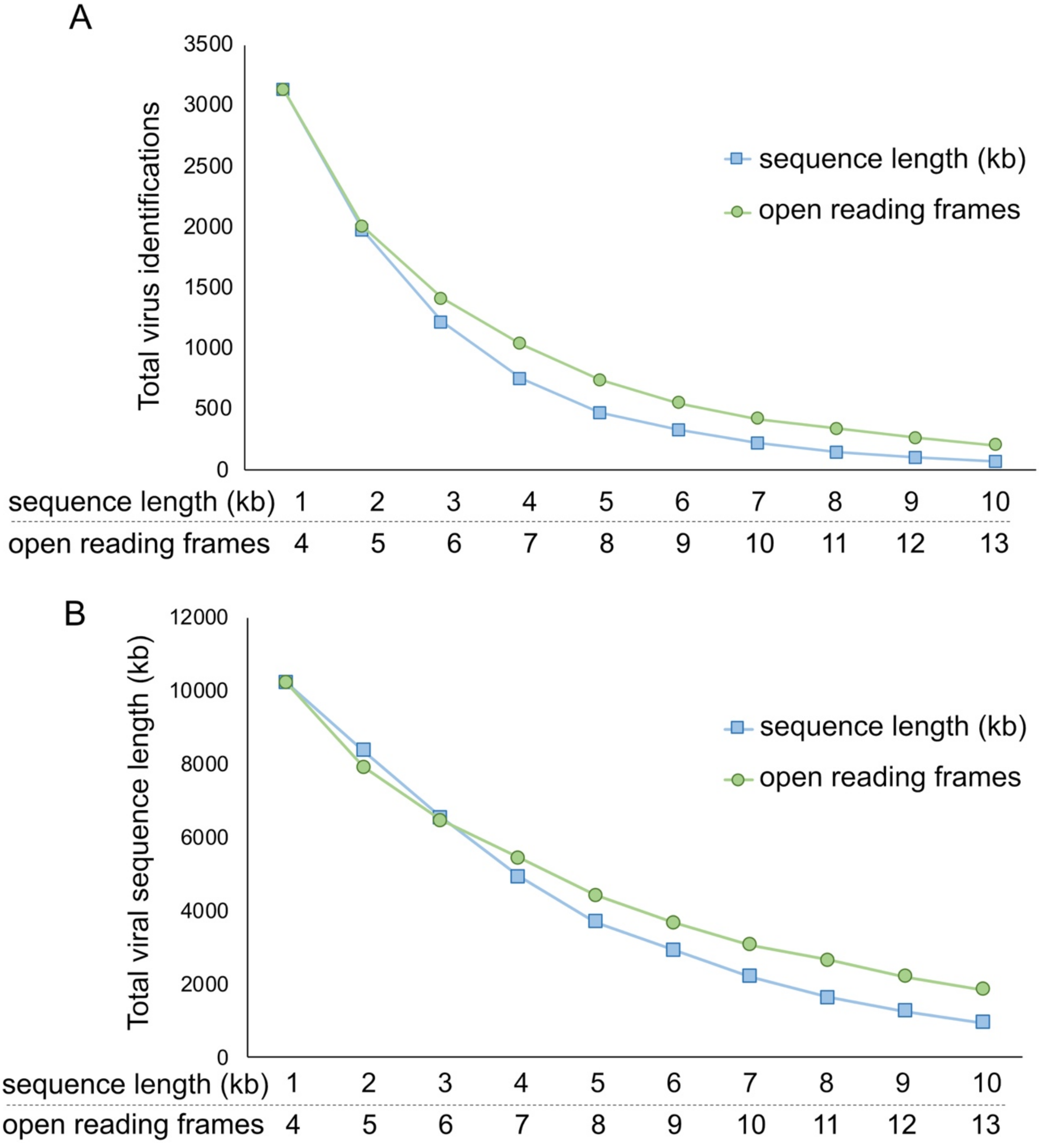


**Supplementary Figure 3. Comparison of limiting to sequence length or open reading frames.** VIBRANT was used to predict viruses from an estuary virome and set to limit to either scaffold length or total encoded open reading frames. The (A) total virus identifications and (B) total viral sequence length were compared to show that limiting to open reading frames will typically yield more data.
